# UniBind: maps of high-confidence direct TF-DNA interactions across nine species

**DOI:** 10.1101/2020.11.17.384578

**Authors:** Rafael Riudavets Puig, Paul Boddie, Aziz Khan, Jaime Abraham Castro-Mondragon, Anthony Mathelier

## Abstract

Transcription factors (TFs) bind specifically to TF binding sites (TFBSs) at cis-regulatory regions to control transcription. Hence, it is critical to locate these TF-DNA interactions to understand transcriptional regulation. The availability of datasets generated by chromatin immunoprecipitation followed by sequencing (ChIP-seq) empowers our efforts to predict the specific locations of TFBSs with greater confidence than previously possible by fusing computational and experimental approaches. In this work, we processed ~10,000 public ChIP-seq datasets from nine species to provide high-quality TFBS predictions. After quality control, it culminated with the prediction of ~56 million TFBSs with experimental and computational evidence for direct TF-DNA interactions for 644 TFs in >1,000 cell lines and tissues. These TFBSs were used to predict >198,000 cis-regulatory modules representing clusters of binding events in the corresponding genomes. The high-quality of the TFBSs was reinforced by their evolutionary conservation, enrichment at active cis-regulatory regions, and capacity to predict combinatorial binding of TFs. Further, we confirmed that the cell type and tissue specificity of enhancer activity was correlated with the number of TFs with binding sites predicted in these regions. All the data is provided to the community through the UniBind database that can be accessed through its web-interface (https://unibind.uio.no/), a dedicated RESTful API, and as genomic tracks. Finally, we provide an enrichment tool, available as a web-service and an R package, for users to find TFs with enriched TFBSs in a set of provided genomic regions. UniBind is the first resource of its kind, providing the largest collection of high-confidence direct TF-DNA interactions in nine species.

## INTRODUCTION

The regulation of gene expression is a complex process involving several biological mechanisms. The first step of the regulatory process controls where, when, and at which intensity RNAs are transcribed from their DNA template. This level of transcriptional regulation is mainly coordinated by transcription factors (TFs), which are DNA-binding proteins that recognize and bind short DNA sequences - their TF binding sites (TFBSs) [1]. TFs are known to co-operate through their combined binding at cis-regulatory regions proximal (promoters) or distal (enhancers or silencers) to the genes they regulate. These regions usually correspond to genomic locations locally dense in TFBSs, which are often referred to as cis-regulatory modules (CRMs), and act as genetic switches to ensure appropriate gene expression [2].

The most popular experimental assay to detect TF-DNA interactions in vivo is chromatin immunoprecipitation followed by sequencing (ChIP-seq) [3]. After mapping the reads generated by ChIP-seq to the genome of interest, the computational analysis aims at identifying genomic regions enriched for mapped reads when compared to a control. The identified genomic locations are known as ChIP-seq peaks. TF ChIP-seq peaks usually span a few hundred base pairs. They derive from direct and indirect TF-DNA interactions [4], where the latter can emerge from protein-protein interactions between the ChIP’ed TF and another protein binding the DNA. Moreover, ChIP-seq peaks could also derive from non-specific binding of the TF to the DNA and noise/bias/artifacts. Several repositories store ChIP-seq peaks [5–9] and are freely available to the community. Nevertheless, these resources do not provide precise locations of the underlying direct TF-DNA interactions.

The TFBSs recognized by a TF are short (~10bp-long) and degenerate sequences that can be modeled computationally for further predictions. The most widely used computational representations of TFBSs for a given TF are position weight matrices (PWMs), which summarize the probability of observing each nucleotide at each position within a set of observed TFBSs. Such computational models have recurrently been used to predict TFBSs in DNA sequences. For instance, one can apply PWMs to predict TFBSs in open chromatin regions (e.g. derived from DNase-seq or ATAC-seq [10–12]) or TF ChIP-seq peaks [13–15].

Previous efforts used PWMs to predict TFBSs within ChIP-seq peaks and made the predictions freely available [15–17]. These resources are specific to one or two species. A substantial limitation of the underlying computational approach is that it relies on the same pre-defined score threshold for all PWMs. Moreover, they do not fully exploit the ChIP-seq peak information such as the enrichment for the TF canonical binding motif close to the ChIP-seq peak summit - where most of the reads align [18]. To address these limitations, we recently developed the ChIP-eat software to specifically delineate direct TF-DNA interactions in ChIP-seq peaks and separate them from indirect or non-specific binding and ChIP-seq artifacts [14]. Briefly, ChIP-eat combines both computational (high PWM score) and experimental (centrality to ChIP-seq peak summit) evidence to find high-confidence direct TF-DNA interactions in a ChIP-seq experiment-specific manner. ChIP-eat was initially applied to 1,983 ChIP-seq peak datasets for 232 human TFs to provide a map of direct TF-DNA interactions in the human genome, which contained >8 million TFBSs stored in the UniBind database [14]. This collection of human TFBSs was proven useful to analyze cis-regulatory alterations in cancers [19–21] and other complex diseases [22,23].

In this report, we describe the update of the UniBind database, which now stores >72 million direct TF-DNA interactions predicted using an updated ChIP-eat pipeline on ~10,000 ChIP-seq peak datasets from nine species: *Arabidopsis thaliana, Caenorhabditis elegans, Danio rerio*, *Drosophila melanogaster, Homo sapiens, Mus musculus, Rattus norvegicus, Saccharomyces cerevisiae*, and *Schizosaccharomyces pombe.* After quality control, we provide the community with a robust collection of ~56 million TFBSs for 644 TFs in 1,097 cell lines and tissues and >198,000 cis-regulatory modules. A functional inspection of these TFBSs and CRMs highlighted that they are evolutionarily conserved and enriched at active cis-regulatory regions. Furthermore, we showed that this unique collection of TFBSs can predict TF binding combinatorics at cis-regulatory regions. Finally, we confirmed that a lower number of TFs binding at enhancers was associated with higher cell type and tissue specificity for these enhancers and vice-versa. The UniBind database is freely available online (https://unibind.uio.no/), through a programmatic RESTful API (https://unibind.uio.no/api/), and via genomic tracks (https://unibind.uio.no/genome-tracks/). Finally, it is accompanied with an enrichment tool to predict TFs with an enrichment of TFBSs in user-provided genomic regions (https://unibind.uio.no/enrichment/).

## RESULTS

### Maps of direct TF-DNA interactions across nine species

#### Prediction of direct TF-DNA interactions

We aimed at providing a collection of direct TF-DNA interactions by combining experimental and computational approaches in several species. We applied an updated version of the ChIP-eat pipeline [14] to ChIP-seq datasets to discriminate high-confidence TFBSs within ChIP-seq peaks from indirect binding events and ChIP-seq noise/artifacts (see Methods). In a nutshell, ChIP-eat uses an entropy-based parameter-free algorithm to automatically define an enrichment zone, which contains TFBSs with high PWM scores and close proximity to ChIP-seq peak summits. These criteria provide computational (high PWM score) and experimental (proximity to peak summit) evidence for direct TF-DNA interactions. This process is carried out in a ChIP-seq dataset specific manner. It first optimizes JASPAR PWMs [24] using the DAMO tool [25] to best discriminate ChIP-seq peaks from random sequences. Next, the optimized PWM is used to detect, for each dataset, the optimal thresholds on the PWM score and distance to the peak summit. These thresholds define the enrichment zone, which highlight direct TF-DNA interactions (see Methods and Gheorghe *et al.* [14] for more details).

We collected ChIP-seq peaks for 11,373 ChIP-seq experiments from ReMap 2018 [26] and GTRD [5] for nine species: *Arabidopsis thaliana*, *Caenorhabditis elegans*, *Danio rerio*, *Drosophila melanogaster, Homo sapiens, Mus musculus, Rattus norvegicus, Saccharomyces cerevisiae* and *Schizosaccharomyces pombe.* For 10,264 datasets, we were able to retrieve a TF binding profile in JASPAR for the ChIP’ed TF. The ChIP-eat pipeline was applied to each ChIP-seq peak dataset - JASPAR PWM pair independently to predict TFBSs. ChIP-eat identified enrichment zones to predict direct TF-DNA interactions in 9,660 datasets. Altogether, this analysis culminated with the prediction of ~72 million TFBSs in ChIP-seq peaks for 841 TFs in 1,317 cell lines and tissues (Supplementary Figure 1; Supplementary Table 1).

We provide these predictions through the UniBind database at https://unibind.uio.no/ (see section UniBind web-application and web-services for details). In the database, the datasets are annotated with information about the ChIP’ed TF (UniProt ID [27]), the cell line or tissue name with ontology IDs from Cellosaurus [28], Cell Line Ontology [29], Experimental Factor Ontology [30], UBERON [31], Cell Ontology [32], and BRENDA [33], and the treatment used, if any.

#### Quality control to establish a robust collection of direct TF-DNA interactions

In UniBind, we aimed to create a robust collection of bona fide direct TF-DNA interactions found in high-quality ChIP-seq peak datasets. This robust collection was obtained by implementing two quality control metrics and only retaining the datasets that satisfy the corresponding criteria. First, we expect high-quality ChIP-seq peak datasets to be enriched for the TF binding motif known to be bound by the ChIP’ed TF. Hence, we filtered out datasets where the DAMO-optimized TF binding motif, which maximizes the discrimination of ChIP-seq peaks from random sequences, was not similar to the expected canonical motif (see Methods). Second, we expect the ChIP-seq peaks to be enriched for TFBSs close to their summits. Hence, we filtered out the datasets where the predicted direct TF-DNA interactions did not show a significant enrichment around the summits (see Methods). While we provide the complete set of TFBSs predicted by ChIP-eat in the permissive collection to the community, we specifically contribute with the robust collection of quality-controlled direct TF-DNA interactions in high-quality ChIP-seq peak datasets.

After applying the quality-control filters, the robust collection of UniBind culminates with ~56 million TFBSs obtained from 6,904 ChIP-seq peak datasets, all species combined (Figure 1A; Supplementary Table 2). No ChIP-seq peak dataset from *S. pombe* passed the quality-control criteria. The TFBSs in the robust collection are associated with 644 distinct TFs ChIP’ed in 1,097 cell lines and tissues (Figure 1A; Supplementary Table 1). We found that the predicted TFBSs covered between 0.04% and 6.06% of the genome of their respective organism (Figure 1B). For example, human and mouse TFBSs covered 6.06% and 5.39% of the genomes, respectively (Figure 1B). Of course, these numbers are somehow a reflection of the number of ChIP-seq experiments available in the corresponding species (Supplementary Figure 2).

**Figure 1.**
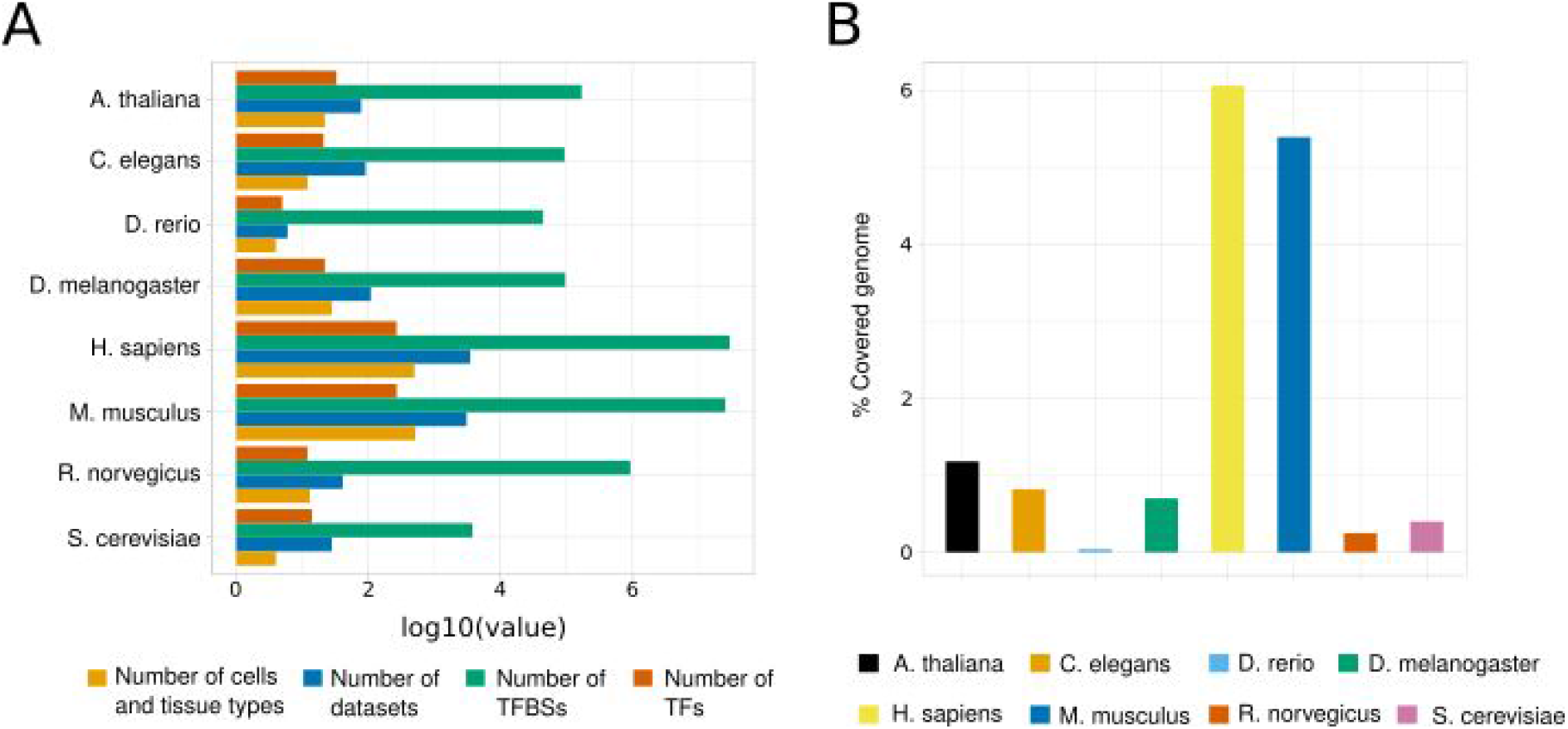
Overview of the UniBind robust collection. (**A**) Barplots showing the number of TFs (dark orange), TFBSs (green), datasets (blue), and cell and tissue types (light orange) stored in the robust collection of UniBind for each analyzed species. All numbers are provided after transformation using the log_10_ function. (**B**) Distribution of the percentages of the genomes covered by robust TFBSs in each species (one color per species, see legend).

Since TFs are known to regulate transcription cooperatively through closely spaced TFBSs [2], we aimed to identify cis-regulatory modules (CRMs) corresponding to clusters of TFBSs. Specifically, we used CREAM [34,35] to locate DNA segments locally enriched for UniBind TFBSs, which culminated with the predictions of >198,000 CRMs (Supplementary Table 2).

To summarize, we provide a collection of TFBSs with both experimental and computational evidence of direct TF-DNA interactions in quality-controlled ChIP-seq peak datasets. Hereafter, the complete collection of unfiltered TFBS predictions is referred to as the “permissive” collection, while the filtered, high-quality TF-DNA interactions are referred to as the “robust” collection.

### Support for the functional relevance of the TFBSs in the robust collection of UniBind

To further confirm the high-quality of the identified TFBSs in the robust collection of UniBind, we sought to provide support for their biological relevance. Hence, the analyses performed below were applied to the robust collection of TFBSs.

#### Human and mouse TFBSs are evolutionarily conserved

We hypothesized that functionally relevant TFBSs should be enriched for evolutionary conservation. Indeed, conservation of DNA segments through evolution represents a hallmark of functional importance [36]. We considered evolutionary conservation scores in the human and mouse genomes computed by the PhyloP [37] and PhastCons [36] methods from the PHAST package [36]. Specifically, we investigated the average conservation of 2 kilobases (kb) DNA regions centered around the TFBS mid-points. For both human and mouse, we noticed that evolutionary conservation gradually increased when the distance to the TFBSs decreased, with sharp peaks of higher conservation at the TFBSs (Figure 2). Increased evolutionary conservation was similarly observed at CRMs (Supplementary Figure 3). The signal was consistently found when considering multiple alignments of 19 (phyloP20way and phastCons20way, Figure 2A) or 99 vertebrate genomes (phyloP100way and phastCons100way, Figure 2A) to the human genome and 59 vertebrate genomes (phastCons60way and phyloP60way, Figure 2B) to the mouse genome. The evolutionary conservation of TFBSs is not expected by chance as no conservation was found when randomly shuffling the positions of the TFBSs in the human and mouse genomes (Figure 2, grey lines). The acute increase of evolutionary conservation scores right at the TFBS locations reinforce the biological relevance of the direct TF-DNA interactions stored in UniBind. This conservation pointed to the potential functional role of the TFBSs in transcriptional regulation of gene expression.

**Figure 2.**
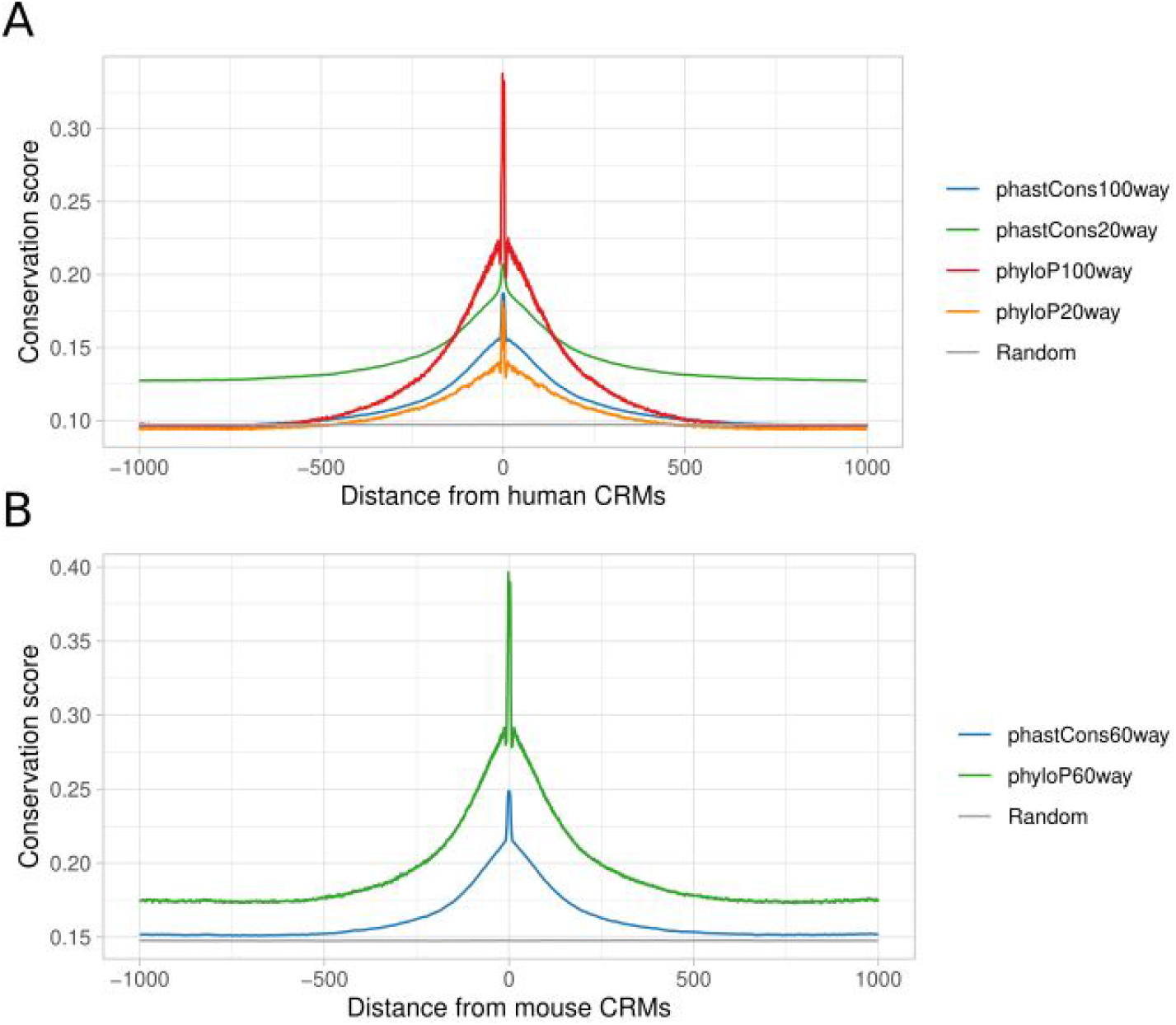
Evolutionary conservation of human and mouse TFBSs in the robust collection. Distributions of the average base-pair evolutionary conservation scores (phyloP and phastCons scores using multi-species genome alignments, see legends) at regions centered around human (**A**) and mouse (**B**) TFBSs from the robust collection. Random expectation (grey lines) was obtained by shuffling the original TFBS locations and obtaining the conservation score of the regions obtained.

#### UniBind TFBSs are enriched at active promoters and enhancers

Next, we sought support for the biological relevance of the TFBSs by assessing their overlap with cis-regulatory regions that are active in different cell types and tissues. We started by mapping out the distribution of the TFBSs with respect to promoter regions, 5’ and 3’ UTRs, exons, introns, regions downstream of genes, and distal intergenic regions (Figure 3, top bar for each species). These distributions were compared to random expectations obtained by shuffling the TFBS positions along the corresponding genomes (Figure 3, bottom bar for each species). By comparing the observed and expected distributions, we noticed that TFBSs were prominently found in promoter regions (<1kb upstream of transcription start sites). The enrichment for TFBSs in promoter regions was further confirmed by (i) OLOGRAM [38], which uses a Monte Carlo simulation approach and a negative binomial model to compute the significance of overlap between two sets of genomic regions (Supplementary Figures 4-9), and (ii) *bedtools reldist* [39], which computes the relative distances between the TFBSs and the genomic regions considered following [40] (Supplementary Figures 10-17). Nevertheless, considering the distribution of TFBSs for each TF independently in each species revealed TFs with binding preferences for promoter regions while others prefer intronic or intergenic regions (Supplementary Figures 18-25). The TSS-proximal versus TSS-distal preferences could explain the previously reported short-versus long-range regulatory effects of TFs [41].

**Figure 3.**
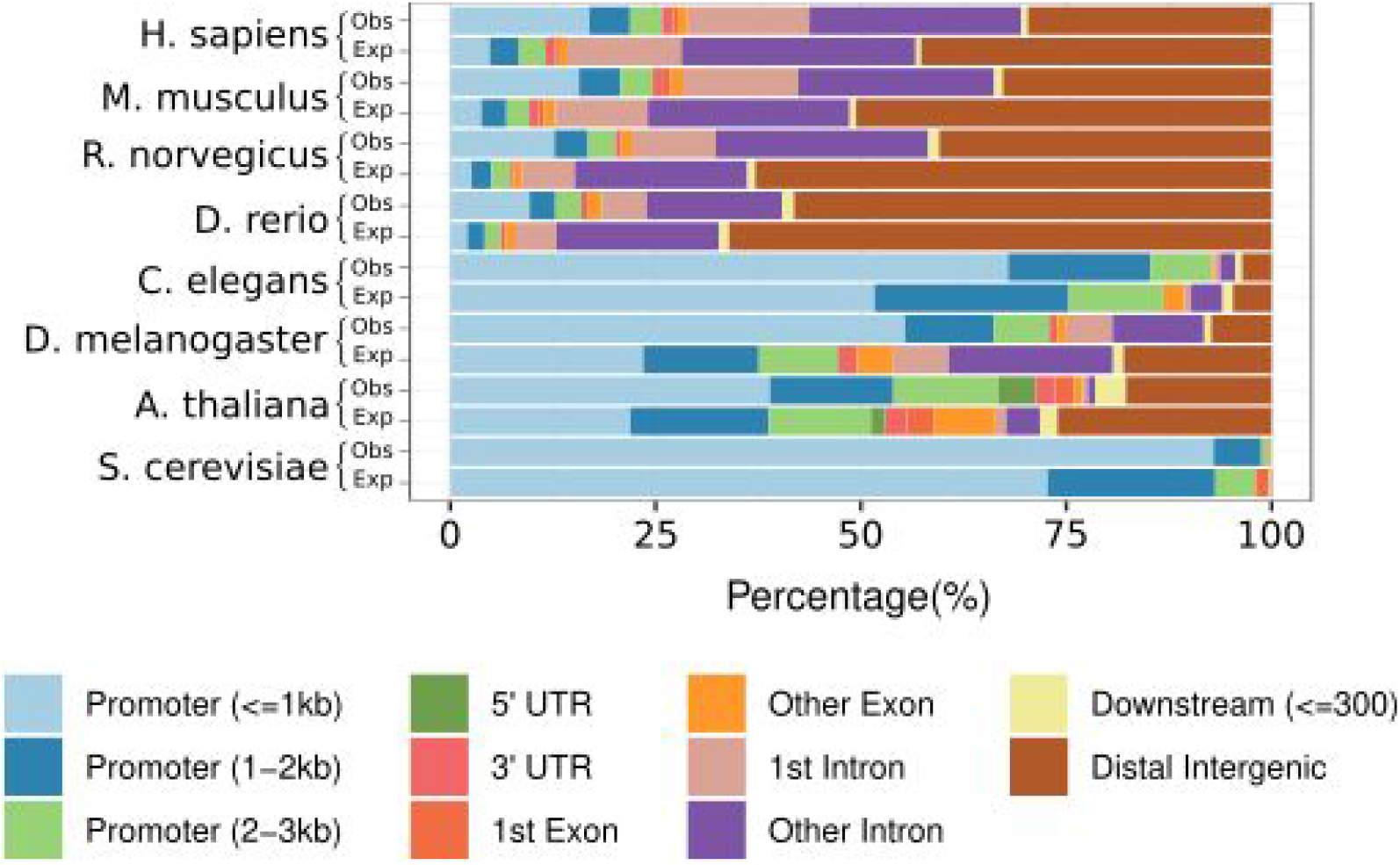
Genomic distribution of TFBSs. Distribution of the proportion of TFBSs from the robust collection overlapping with different types of genomic regions (columns; see legend) across species (rows). For each species, we provide the observed (first lines, denoted Obs) and expected (second lines, denoted Exp) proportions of TFBSs in each type of genomic regions. Expected proportions were estimated by randomly positioning the TFBSs in the corresponding genomes (see Methods).

In the vertebrate species (human, mouse, rat, and zebrafish), the majority of TFBSs lie in introns and distal intergenic regions (Figure 3), which is expected given the large portion of the corresponding genomes covered by these non-coding regions. To confirm the biological function of the TFBSs stored in UniBind, we examined their overlap with active cis-regulatory regions in the mouse and human genomes. We considered the candidate cis-regulatory elements (cCREs) predicted using epigenetic marks by the ENCODE consortium [42]. Specifically, DNase I hypersensitive open chromatin regions were first identified and then overlapped with H3K4me3 and H3K27ac histone modification marks and CTCF ChIP-seq data to predict five types of cCREs with: (1) a promoter-like signature (PLS), (2) an enhancer-like signature proximal (pELS) or (3) distal (dELS) to TSSs, (4) a H3K4me3 signature (DNase-H3K4me3), or (5) a CTCF-only signature [42]. Consistently, we confirmed that UniBind TFBSs were enriched in PLS and ELS cCREs when considering both the OLOGRAM and *bedtools reldist* evaluations of overlap (Figure 4A-B; Supplementary Figures 26-27). The enrichment at regions of active promoter signature is consistent with the genomic distribution observed above. The enrichment at regions harbouring active enhancer signature suggests that the TFBSs are not randomly spaced in the introns and intergenic regions. Furthermore, we confirmed that CRMs were enriched for cCREs with active promoter- or enhancer-like signatures when considering the 105,104 and 73,917 CRMs predicted in human and mouse, respectively (Supplementary Figures 28-30). Figure 4C shows an example of the UCSC Genome Browser [43] at the LDLR gene locus where we observe the overlap between UniBind TFBSs, CRMs, and cCREs.

**Figure 4.**
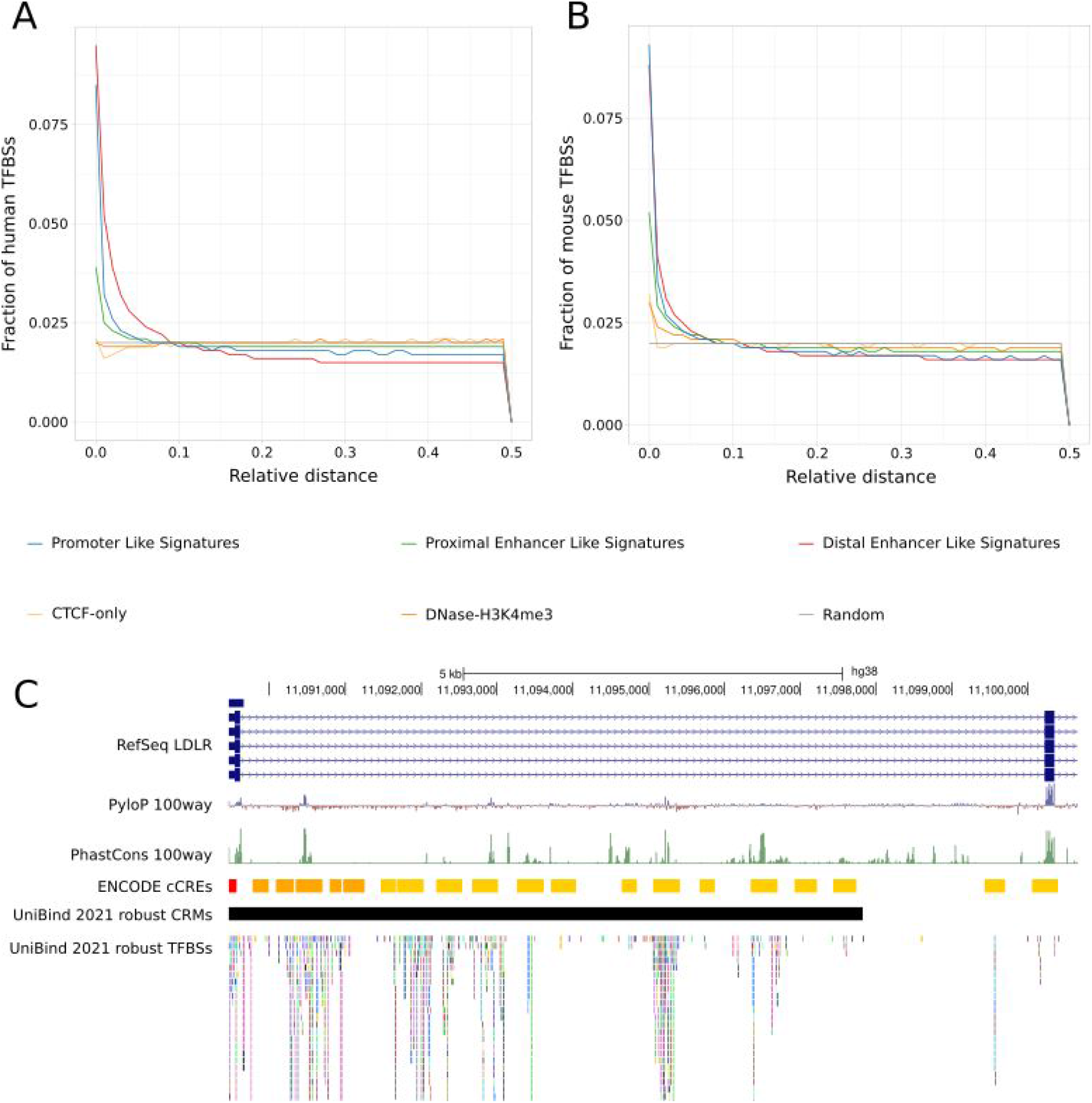
Analysis of the overlap of TFBSs with respect to active cis-regulatory regions in human and mouse. (**A-B**) Fraction of TFBSs in the UniBind robust collection (y-axis) with respect to increasing relative distances (x-axis) from ENCODE candidate cis-regulatory regions (cCREs) computed using the *bedtools reldist* command for human (A) and mouse (B). When two genomic tracks are not spatially related, one expects the fraction of relative distance distribution to be uniform. (**C**) Genomic tracks from the UCSC Genome Browser at the LDLR gene locus (from start to first coding exon) providing information about PhyloP and PhastCons evolutionary conservation scores and the locations of ENCODE cCREs, UniBind CRMs, and UniBind TFBSs from the robust collection. Colors in the ENCODE cCREs track indicate: Promoter like signature (red), proximal enhancer like signature (orange), and distal enhancer like signature (yellow).

Together, these results highlight the biological relevance of the UniBind TFBSs and CRMs for transcriptional regulation via their association with active promoters and enhancers in human and mouse.

#### Specificity of enhancer activity in cell types and tissues correlates with binding TF composition

We further investigated how the number of TF binding events at enhancers could be related to their regulatory effects. We considered enhancers that were identified through the capture of bidirectional transcription of enhancer RNAs (eRNAs) at their boundaries using Cap Analysis of Gene Expression (CAGE) in 1,829 human libraries [44]. Cell type and tissue specificity was assessed by considering the amount of eRNAs captured by CAGE across the libraries [44]. We overlapped the UniBind TFBSs with the CAGE-derived enhancers and assessed the relationship between the expression specificity of the enhancers and the number of TFs with binding sites in these enhancers. We observed that cell type / tissue specific enhancers tend to harbour a lower number binding TFs, while more ubiquitously active enhancers tend to harbour a higher number of binding TFs (Figure 5; Supplementary Figure 31). The correlation between the number of binding TFs and cell type / tissue expression specificity of enhancers is inline with previous observations showing an association between the number of TFBSs and the combinatorics of TFs at promoters and enhancers with enhancer activity strength and specificity [45–47]. Altogether, these observations underline the importance of TF cooperation for cis-regulatory activity.

**Figure 5.**
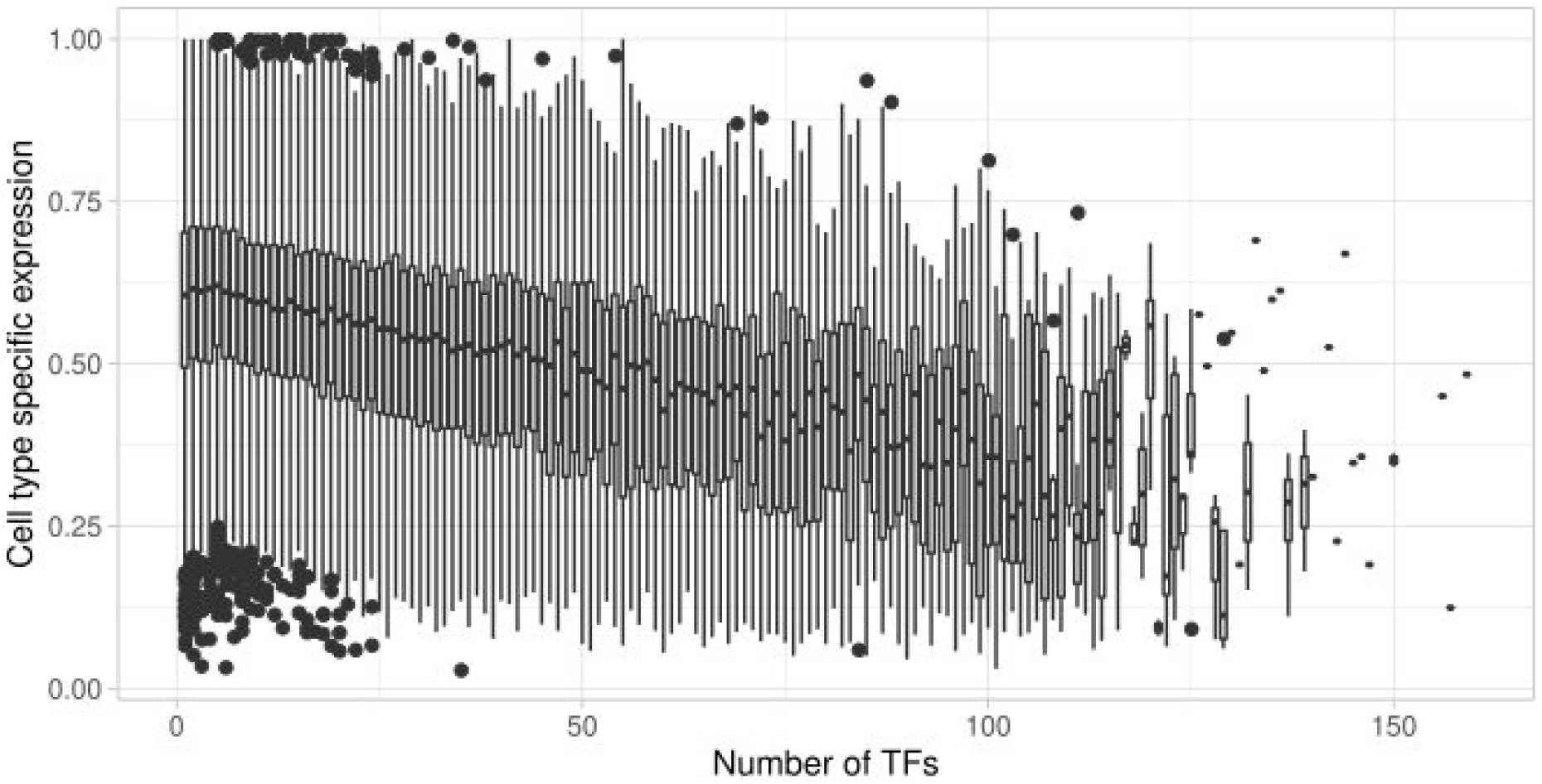
Correlation between enhancer activity and TF binding. For each enhancer predicted using Cap Analysis of Gene Expression (CAGE) by the FANTOM5 consortium, we computed the number of TFs with overlapping TFBSs in the robust collection of UniBind (x-axis). The figure provides, for each value of the number of TFs found to bind in enhancers, a boxplot of the distribution of cell type specific activity of these enhancers. The expression measures were derived from CAGE (capturing enhancer RNA expression). The specificity of activity (y-axis) is provided within the [0; 1] range with 0 representing ubiquitous enhancer activity and 1 exclusive expression activity.

#### UniBind TFBSs reveal TF binding combinatorics at cis-regulatory regions

We explored the capacity of UniBind TFBSs to further pinpoint relevant TF binding combinatorics at cis-regulatory regions. As a case study, we examined the direct TF-DNA interactions stored in UniBind and derived from ChIP-seq experiments in the untreated MCF7 cell line. This cell line is representative of estrogen positive (ER+) invasive ductal breast carcinoma, which is known to be mainly driven by the combined activity of the TFs ESR1, GATA3, and FOXA1 [48]. We extended the genomic locations of UniBind TFBSs predicted in MCF7 by 50bp on each side and intersected these regions between each pair of MCF7 TFBS datasets using the Intervene tool [49]. Next, we computed the fractions of overlap for each pair and calculated the pairwise Pearson correlation coefficients of the fractions of overlap between all pairs of datasets. A high pairwise correlation coefficient between two datasets indicates that the underlying TFBS regions are co-localizing. Hierarchical clustering of the pairwise correlation coefficient revealed 4 main clusters (Figure 6). As expected, we observed high correlations between datasets for the same TF (e.g. red cluster in Figure 6 with exclusively CTCF TFBSs). The largest cluster (Figure 6, green) was mainly composed of TFBSs from ESR1, FOXA1, and GATA3. Co-localization of binding events for these TFs confirm the potential of UniBind TFBSs to highlight TFs known to cooperate at cis-regulatory regions. The second largest cluster (Figure 6, blue) contained TFBSs for E2F1, NRF1, MAX, MYC, ELK1, ELF1, GABPA, EGR1, and SRF. Among these TFs, MAX and MYC as well as ELK1 and SRF are known to dimerize to bind DNA. Finally, the purple cluster was composed of JUN and FOS TFBSs, known to bind DNA as a dimer to form the AP1 complex. This case study exemplifies how UniBind TFBSs can be used to derive biologically relevant information about TF binding combinatorics.

**Figure 6.**
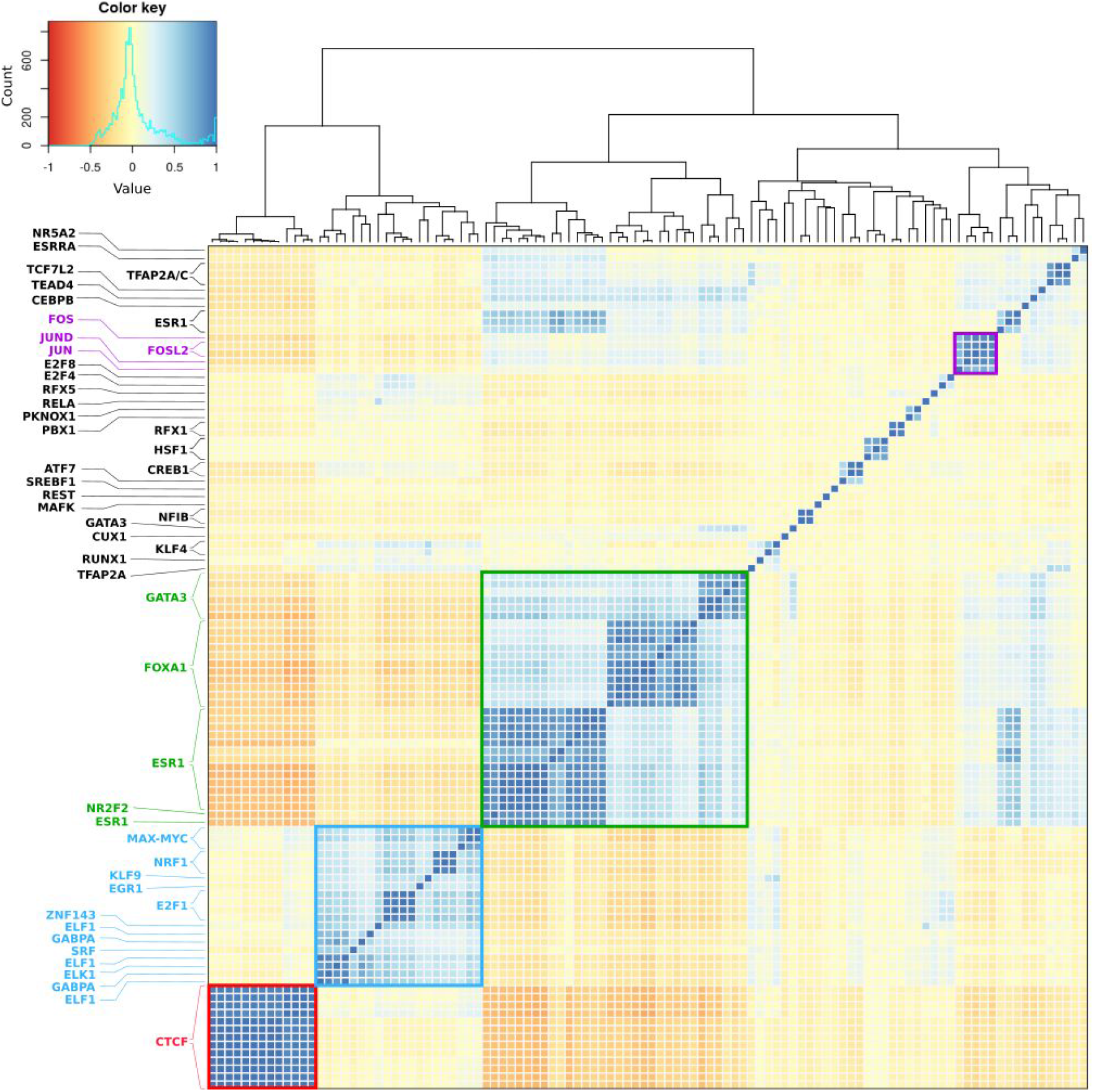
TF combinatorial binding in invasive breast ductal carcinoma. Hierarchical clustering of the pairwise Pearson correlation coefficient between all TFBSs from untreated MCF7 cells from the robust collection of UniBind. Different clusters and their respective TFs are coloured in red, blue, green and purple. In the heatmap, blue colors indicate a higher positive correlation coefficient between datasets, while red colors indicate an anticorrelation (see legend).

Altogether, the assessments of the functional and biological relevance to study transcriptional regulation outlined here support, a posteriori, the high-quality of the direct TF-DNA interactions stored in the robust collection of UniBind.

### UniBind web-application and web-services

#### Accessing and exploring UniBind data

All the direct TF-DNA interactions from the permissive and robust collections are freely available through the UniBind web-application at https://unibind.uio.no. The predictions come with metadata about the associated ChIP-seq experiments and external links to useful resources such as ReMap [26], GeneCards [50], and GEO [51]. Users can search and explore the data through the user-friendly web-interface. The web-application provides a search interface for users to filter the datasets using the metadata fields and search results are downloadable as a metadata table as well as FASTA and BED files for the TFBSs. To improve the search ability of the data, the search engine supports gene synonyms when searching for TFs. All data can be downloaded for individual datasets as well as through bulk download links per species or collection. In addition, we developed a RESTful API (https://unibind.uio.no/api/) to allow programmatic access to the stored data from any programming language. Finally, we built genome track hubs that are easily visualized through the UCSC [52] and Ensembl [53] genome browsers. The track hubs can be accessed through the UniBind web-application (https://unibind.uio.no/genome-tracks/) as well as through the public track hubs at UCSC [52] and the track hub registry (https://trackhubregistry.org/).

#### TFBS sets enrichment application tool

A regular task when studying transcriptional regulation is to find TFs that are the most likely to control the activity of a set of cis-regulatory regions. Classical strategies rely on the prediction of enriched potential TFBSs for a set of TFs derived from either ChIP-seq peaks datasets [54–57] or PWM predictions [56,58]. As UniBind stores TFBSs with both ChIP-seq and PWM evidence of direct TF-DNA interactions, one can rely on this resource to infer the TFs likely to bind a set of cis-regulatory regions. The method consists in computing the enrichment for specific TFBS sets in given DNA regions compared to background regions. We provide a web-service (and the underlying source code) to perform this TFBS dataset enrichment analysis to the users at https://unibind.uio.no/enrichment/ (Figure 7A). The enrichment computation relies on the Locus Overlap Analysis (LOLA) tool [59]. The enrichment tool provides three different types of enrichment analyses: (1) using a provided universe of potentially bound regions; (2) comparing enrichment with another set of genomic regions to perform differential enrichment; or (3) comparing the enrichment to all TFBSs stored in UniBind as a universe (Figure 7A).

**Figure 7.**
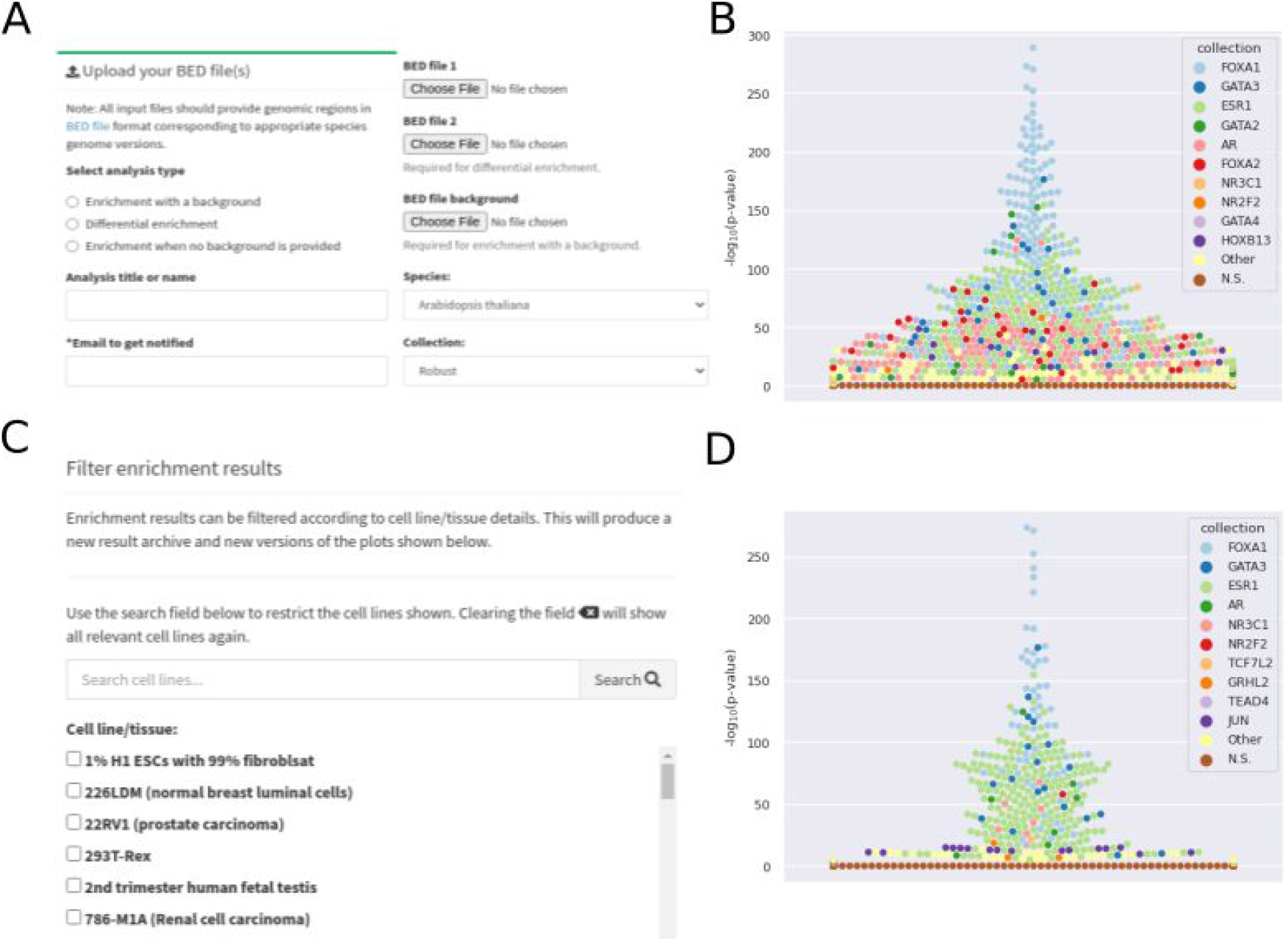
The UniBind TFBS set enrichment tool. (**A**) The UniBind enrichment web-application allows users to select the enrichment analysis type, set a title, provide an email address for notification upon completion of the analysis, upload of the required input files based on the enrichment analysis type, and select the species and collection to compute the enrichment. (**B**) Enrichment results shown as swarm plots of the -log_10_(p-values) (Fisher exact tests; see Methods). Each point corresponds to a TFBS set for a given TF in a given ChIP-seq experiment. Distinct colors are assigned to the top 10 TFs with at least one TFBS set enriched (see legend). (**C**) The enrichment results can be further explored by restricting the output to TFBS sets obtained in specific cell lines and tissues, which can be searched by keywords and selected. (**D**) Swarm plot similar to (B) but restricted to TFBS sets obtained from breast-related tissues and cell lines.

As a case example, we applied the enrichment tool to genomic regions surrounding CpGs found to be demethylated in ER+ breast cancer patients [60]. As a background set, we used all CpG probes from the Illumina Infinium HumanMethylation450 microarray. The demethylated CpG regions in ER+ patients were predicted to be bound by FOXA1, GATA2, GATA3, ESR1 and AR (top 5 TFs, Figure 7B). The enrichment of these TFs is in line with the known ER+ TF drivers [60]. Further, the enrichment tool allows users to filter the results by restricting the search to TFBS datasets derived from specific cell lines / tissues. In our case study, limiting to breast-related cell types and tissues highlights FOXA1, GATA3, and ESR1 with the most enriched TFBS sets (Figure 7C-D), which is in agreement with the driving role of this TFs in ER+ carcinogenesis.

## CONCLUSIONS AND PERSPECTIVES

Through the uniform processing of >10,000 ChIP-seq peak datasets, we provide maps of direct TF-DNA interactions in nine species. Altogether, this process culminated with the prediction of >72 million TFBSs, ~56 million of which passed stringent QC criteria to compose the robust collection of direct TF-DNA interactions in UniBind. The robust collection is associated with 644 distinct TFs from 6,904 ChIP-seq datasets derived from 1,097 cell lines and tissues. Functional assessments of the robust collection of TFBSs through evolutionary conservation and strong overlap with active promoters and enhancers in human and mouse highlighted the high-quality and biological relevance of the collection. Further, we showed that the TFBSs can provide insights into enhancer activity and TF binding combinatorics at cis-regulatory regions. Previous works combined with the results outlined here underline the functional relevance of analyzing TFBS combinatorics at cis-regulatory elements to shed light on the molecular mechanisms underlying transcriptional regulation.

We provide this resource freely to the community through a dedicated web-application, a RESTful API, and genome tracks for the UCSC and Ensembl genome browsers. Finally, TFBS dataset enrichment analyses can be performed through an online web-service and a stand-alone tool to predict the TFs acting upon a set of genomic regions.

The TFBS predictions provided in the current version of UniBind were obtained using PWMs as computational models. In the original version dedicated to human [14], we provided predictions obtained from four different computational models: PWMs, binding energy models [61], transcription factor flexible models (TFFMs) [62], and DNA shape-based models (DNAshapedTFBS models) [63]. The ChIP-eat pipeline is agnostic to the computational model used to predict the enrichment zone with high computational and experimental evidence of direct TF-DNA interactions. Hence, we foresee that more sophisticated models than PWMs could be used to predict TFBSs to be stored in UniBind in the future, should they become extensively used by the community.

UniBind relies on the availability of ChIP-seq peak datasets made available to the community. The current release relies on the ReMap and GTRD databases. These databases were selected as they (1) encompass a large part of the publicly available ChIP-seq experiments for several species, (2) process ChIP-seq data uniformly, and (3) are regularly updated and under active maintenance. UniBind will be updated on a regular basis, as soon as new ChIP-seq datasets become available in ReMap and GTRD. Moreover, we are open to including other ChIP-seq peak resources that fulfill the criteria described above (e.g. repositories specialized in some species or taxa) for the upcoming updates of UniBind.

## MATERIAL AND METHODS

### ChIP-seq peak datasets and TF binding profiles

A total of 11,373 ChIP-based datasets with peaks predicted by MACS [64] were retrieved from the ReMap (2018 version) [26] and GTRD [5] databases. ReMap datasets were the same as the ones used in the previous UniBind release and were reprocessed with new JASPAR PWMs. Note that some datasets were obtained using the ChIP-exo or ChIP-nexus protocols; we refer to ChIP-seq datasets as a whole in this manuscript for simplicity.

ChIP-seq peak datasets were associated with JASPAR (version 2020) [24] TF binding profiles (provided as position frequency matrices, PFMs) whenever possible. Specifically, we used the HGNC gene symbols to search the collection of JASPAR TF binding profiles in the same taxonomic group as the ChIP’ed TF. For the datasets where no TF binding profile was found, we used the *mygene* bioconductor package [65] to obtain all possible gene synonyms and used the synonyms to search for JASPAR TF binding profiles. Altogether, JASPAR PFMs were assigned to 10,264 datasets out of 11,373. Note that some ChIP-seq peak datasets stored in ReMap and GTRD are not associated with TFs but general transcriptional regulators (e.g. EP300, RAD21, SMC4), so no PFM in JASPAR could be assigned; for some TFs, no PFM was available in JASPAR.

### Genome assemblies

The genome assemblies used for each species were: *hg38 (H. sapiens), mm10 (M. musculus), Rnor_6.0 (R. norvegicus), WBcel235 (C. elegans), dm6 (D. melanogasteŕ), GRCz11 (D. rerio), TAIR10 (A. thaliana), R64-1-1 (S. cerevisiae),* and *ASM294v2 (S. pombe).*

### Identification of direct TF-DNA interactions

We applied the ChIP-eat pipeline (https://bitbucket.org/CBGR/chip-eat/src/master/) to each ChIP-seq peak dataset independently, following a similar method as described previously [14]. Compared to the original version of ChIP-eat [14], we made the two following modifications: (1) we used DAMO [25] (version 1.0.1) with default parameters to optimize the PWMs in a dataset-specific manner; (2) once the thresholds (on the distance to peak summits and PWM score) defining the enrichment were predicted, we rescanned the peaks with the DAMO-optimized PWMs and kept the best hit (highest PWM score) per peak that fall within the enrichment zone, if any.

### Quality control metrics for the robust collection

TFBSs in the permissive collection were filtered using two quality control (QC) metrics. (1) To ensure similarity between the DAMO-optimized PFM and the original JASPAR PFM, we only kept in the robust collection the datasets providing a TOMTOM (version 4.11.4) [66] similarity p-value strictly below 0.05. This QC metric ensures that the canonical motif known to be recognized by the ChIP’ed TF is enriched in the ChIP-seq peaks. (2) To ensure a strong enrichment for direct TF-DNA interactions in the vicinity of the peak summits, we computed a centrality enrichment following the method described in CentriMo [67]. Only TFBS datasets with a centrality p-value < 0.05 were kept in the robust collection. This QC metric ensures that TFBSs are enriched in the vicinity of the peak summits overall in the ChIP-seq peaks considered (some of which are not predicted to contain a direct TF-DNA interactions / TFBS).

### Cis-regulatory modules

For each species, we considered unique locations of permissive and robust TFBSs separately and used CREAM [35] with default parameters to compute cis-regulatory modules.

### Random positioning of TFBSs

The random distribution of TFBSs was obtained by shuffling the original unique TFBS coordinates along the genomes using the *shuffle* subcommand of the *bedtools* (version 2.25.0) [39] with the *-chrom* option to keep the same number of TFBSs per chromosome.

### Evolutionary conservation

The evolutionary conservation scores were retrieved from the UCSC genome browser data portal as bigWig files for the human and mouse genomes. Specifically, we downloaded the bigWig files corresponding to the tracks phastCons100way, phastCons20way, phyloP100way, and phyloP20way for human and phastCons60way and phyloP60way for mouse. We considered unique locations of human and mouse TFBSs from the robust collection and the average conservation scores in 2kb regions centered around the TFBS mid-points were computed using the *agg* subcommand of *bwtool* (version 1.0) [68]. The same strategy was applied to the random positions of TFBSs and the CRMs.

### Genomic distributions

For each species, the genomic coordinates of all TFBSs were retrieved and duplicate coordinates (from multiple ChIP-seq experiments) were filtered out to conserve only unique genomic locations. The distributions of these unique TFBS positions with respect to promoters, 5’ and 3’ UTRs, exons, introns, regions downstream of genes, and intergenic regions were obtained using the ChIPseeker Bioconductor package (version 1.20.0) [69]. We used the following genome annotations with ChIPseeker: TxDb.Athaliana.BioMart.plantsmart28 *(A. thaliana),* TxDb.Celegans.UCSC.ce11.refGene *(C. elegans),* TxDb.Drerio.UCSC.danRer11.refGene *(D. rerio),* TxDb.Dmelanogaster.UCSC.dm6.ensGene (*D. melanogasteŕ),* TxDb.Hsapiens.UCSC.hg38.knownGene *(H. sapiens),* and TxDb.Mmusculus.UCSC.mm10.knownGene *(M. musculus).* The genome annotations for *R. norvegicus* and *S. cerevisiae* were built from GTF files obtained from Ensembl by using the *makeTxDbFromGFF* function from the *GenomicFeatures* Bioconductor package [70] (version 1.36.4). The same methodology was applied to the random distribution of TFBSs.

The enrichment for the unique TFBS positions at the different genomic features was computed using the OLOGRAM function of the *gtftk* package (version 1.2.1) [38,71]. Note that no result is provided for *H. sapiens* and *M. musculus* as OLOGRAM did not manage to complete the computations.

### Relative distances and enrichment with candidate cis-regulatory elements (cCREs)

The genomic coordinates of human and mouse cCREs predicted by ENCODE were retrieved as BED files from the SCREEN web-portal at https://screen.encodeproject.org/.

The relative distances between the unique TFBS positions and the ENCODE cCREs were computed using the *reldist* subcommand of the *bedtools* (version 2.25.0). The same methodology was applied to the CRMs and the randomly distributed TFBSs.

The enrichment for the unique TFBS positions at the ENCODE cCREs was computed using the OLOGRAM function of the *gtftk* package (version 1.2.1) [38,71]. The same methodology was applied to the CRMs.

### Cell type and tissue specific enhancer expression

The genomic coordinates (hg19 genome assembly) of the 43,011 permissive enhancers predicted from CAGE experiments [44] were retrieved as BED files from http://enhancer.binf.ku.dk/presets/. Coordinates were converted to the hg38 genome assembly using the UCSC *liftOver* tool [72]. For each TF, we considered unique genomic coordinates and intersected these locations with the enhancer coordinates using the *intersect* subcommand of the *bedtools* (version 2.29.2) using the options *-wa -filenames -C.* The results were used to compute the number of TFs with at least one TFBS overlapping the enhancers.

Enhancer cell type and tissue specific expressions were obtained from Andersson *et al.* [44] and computed as 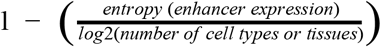. The vector of expression values for each enhancer over cell types or tissues corresponded to the mean of the enhancer expression in each cell type or tissue [44].

### Pairwise correlation computation for TFBS datasets from MCF7

The TFBS datasets associated with the MCF7 cell line were retrieved from the UniBind database using the search functionality of the web-application. Metadata was used to restrict the datasets to the ones where no treatment was introduced in the MCF7 cells. For each dataset, TFBS positions were expanded by 50bp on each side using the *slop* subcommand of the *bedtools* and then merged using the *sort* and *merge* subcommands of the *bedtools*. These genomic regions were used as input to the *pairwise* subcommand of the Intervene tool [49] to compute the fraction of intersections between each pair of datasets. Pairwise Pearson correlation coefficients between the vectors of fraction of intersections between each pair of datasets were computed using Intervene. Hierarchical clustering was obtained through the Intervene Shiny application (https://intervene.shinyapps.io/intervene/) with the *Heatmap.2* function.

### Genome track hubs

Genome track hubs were built following the specifications at https://genome.ucsc.edu/goldenPath/help/hgTrackHubHelp.html.

### Enrichment tool and web-service

The enrichment tool relies on the LOLA Bioconductor package (version 1.14.0) [59] to assess enrichment of overlaps based on Fisher exact tests. For each species, a dedicated LOLA database was built with all the predicted TFBSs and the corresponding metadata informing about cell type / tissue, treatment, and TF name. The databases were generated following the instructions provided at http://databio.org/regiondb and are available as RDS R objects on Zenodo at https://doi.org/10.5281/zenodo.4452896. The web-service is freely available at https://unibind.uio.no/enrichment/ with source code for the standalone software available at https://bitbucket.org/CBGR/unibind_enrichment/.

### UniBind web-application

The UniBind web-application is developed in Python using the model-view-controller framework Django. It uses SQLite to store TFBS metadata and Bootstrap as the frontend template engine. The search function relies on the RESTful API (see below). It allows for searching for gene name synonyms using naming data from the Entrez Gene and SwissProt databases and combining such data with JASPAR matrix profile information to yield a relevant collection of synonyms (source code at https://bitbucket.org/CBGR/synonyms). The source code of the UniBind web-application together with installation instructions are available at https://bitbucket.org/CBGR/unibind.

### RESTful API

The RESTful API is implemented in Python as part of the UniBind web-application using the Django REST Framework. An Apache HTTP server provides access to the application and thus to the API, with the underlying SQLite database system supporting queries constructed by the API implementation to retrieve data requested by users of the API. The available REST API endpoints are “Datasets”, “Cell types”, “Collections”, “Species”, and “Transcription factors”. The API is available at https://unibind.uio.no/api/.

## ACKNOWLEDGEMENTS

We thank Roza Berhanu Lemma for her helpful comments on the manuscript, and Ieva Rauluseviciute and Oriol Fornés for their valuable feedback and testing. We thank Georgios Magklaras, Harold Gutch, and the NCMM IT team for their IT support, and Ingrid Kjelsvik for her administrative support.

## REFERENCES

1. Suter DM. Transcription Factors and DNA Play Hide and Seek. Trends Cell Biol. 2020. doi: 10.1016/j.tcb.2020.03.003

2. Wasserman WW, Sandelin A. Applied bioinformatics for the identification of regulatory elements. Nat Rev Genet. 2004;5: 276–287.

3. Johnson DS, Mortazavi A, Myers RM, Wold B. Genome-Wide Mapping of in Vivo Protein-DNA Interactions. Science. 2007;316: 1497–1502.

4. Furey TS. ChIP-seq and beyond: new and improved methodologies to detect and characterize protein-DNA interactions. Nat Rev Genet. 2012;13: 840–852.

5. Yevshin I, Sharipov R, Kolmykov S, Kondrakhin Y, Kolpakov F. GTRD: a database on gene transcription regulation—2019 update. Nucleic Acids Res. 2018;47: D100–D105.

6. Chèneby J, Ménétrier Z, Mestdagh M, Rosnet T, Douida A, Rhalloussi W, et al. ReMap 2020: a database of regulatory regions from an integrative analysis of Human and Arabidopsis DNA-binding sequencing experiments. Nucleic Acids Res. 2020;48: D180–D188.

7. Mei S, Qin Q, Wu Q, Sun H, Zheng R, Zang C, et al. Cistrome Data Browser: a data portal for ChIP-Seq and chromatin accessibility data in human and mouse. Nucleic Acids Res. 2016;45: D658–D662.

8. Zhou K-R, Liu S, Sun W-J, Zheng L-L, Zhou H, Yang J-H, et al. ChIPBase v2.0: decoding transcriptional regulatory networks of non-coding RNAs and protein-coding genes from ChIP-seq data. Nucleic Acids Res. 2017;45: D43–D50.

9. Chen D, Fu L-Y, Zhang P, Chen M, Kaufmann K. ChIP-Hub: an Integrative Platform for Exploring Plant Regulome. Bioinformatics. bioRxiv; 2019. p. 784.

10. Vierstra J, Lazar J, Sandstrom R, Halow J, Lee K, Bates D, et al. Global reference mapping of human transcription factor footprints. Nature. 2020;583: 729–736.

11. Bentsen M, Goymann P, Schultheis H, Klee K, Petrova A, Wiegandt R, et al. ATAC-seq footprinting unravels kinetics of transcription factor binding during zygotic genome activation. Nat Commun. 2020;11: 4267.

12. Li Z, Schulz MH, Look T, Begemann M, Zenke M, Costa IG. Identification of transcription factor binding sites using ATAC-seq. Genome Biol. 2019;20: 45.

13. Wang J, Zhuang J, Iyer S, Lin X, Whitfield TW, Greven MC, et al. Sequence features and chromatin structure around the genomic regions bound by 119 human transcription factors. Genome Res. 2012;22: 1798–1812.

14. Gheorghe M, Sandve GK, Khan A, Chèneby J, Ballester B, Mathelier A. A map of direct TF-DNA interactions in the human genome. Nucleic Acids Res. 2018;47: e21–e21.

15. Czipa E, Schiller M, Nagy T, Kontra L, Steiner L, Koller J, et al. ChIPSummitDB: a ChIP-seq-based database of human transcription factor binding sites and the topological arrangements of the proteins bound to them. Database. 2020;2020. doi:10.1093/database/baz141

16. Fornes O, Gheorghe M, Richmond PA, Arenillas DJ, Wasserman WW, Mathelier A. MANTA2, update of the Mongo database for the analysis of transcription factor binding site alterations. Sci Data. 2018;5: 180141.

17. Bülow L, Brill Y, Hehl R. AthaMap-assisted transcription factor target gene identification in Arabidopsis thaliana. Database. 2010;2010: baq034.

18. Worsley Hunt R, Mathelier A, del Peso L, Wasserman WW. Improving analysis of transcription factor binding sites within ChIP-Seq data based on topological motif enrichment. BMC Genomics. 2014;15: 472.

19. Singh AK, Talseth-Palmer B, McPhillips M, Lavik LAS, Xavier A, Drabløs F, et al. Targeted sequencing of genes associated with the mismatch repair pathway in patients with endometrial cancer. PLoS One. 2020;15: e0235613.

20. Castro-Mondragon JA, Aure MR, Lingærde OC. Cis-regulatory mutations associate with transcriptional and post-transcriptional deregulation of the gene regulatory program in cancers. bioRxiv. 2020. Available: https://www.biorxiv.org/content/10.1101/2020.06.25.170738v1.abstract

21. Uusi-Mäkelä J, Afyounian E, Tabaro F, Häkkinen T. Chromatin accessibility analysis uncovers regulatory element landscape in prostate cancer progression. bioRxiv. 2020. Available: https://www.biorxiv.org/content/10.1101/2020.09.08.287268v1.abstract

22. Rhead B, Shao X, Quach H, Ghai P, Barcellos LF, Bowcock AM. Global expression and CpG methylation analysis of primary endothelial cells before and after TNFa stimulation reveals gene modules enriched in inflammatory and infectious diseases and associated DMRs. PLoS One. 2020;15: e0230884.

23. Wang X, Goldstein DB. Enhancer Domains Predict Gene Pathogenicity and Inform Gene Discovery in Complex Disease. Am J Hum Genet. 2020;106: 215–233.

24. Fornes O, Castro-Mondragon JA, Khan A, van der Lee R, Zhang X, Richmond PA, et al. JASPAR 2020: update of the open-access database of transcription factor binding profiles. Nucleic Acids Res. 2019;48: D87–D92.

25. Ruan S, Stormo GD. Comparison of discriminative motif optimization using matrix and DNA shape-based models. BMC Bioinformatics. 2018;19: 86.

26. Chèneby J, Gheorghe M, Artufel M, Mathelier A, Ballester B. ReMap 2018: an updated atlas of regulatory regions from an integrative analysis of DNA-binding ChIP-seq experiments. Nucleic Acids Res. 2017;46: D267–D275.

27. The UniProt Consortium. UniProt: the universal protein knowledgebase. Nucleic Acids Res. 2017;45: D158–D169.

28. Bairoch A. The Cellosaurus, a Cell-Line Knowledge Resource. J Biomol Tech. 2018;29: 25–38.

29. Sarntivijai S, Lin Y, Xiang Z, Meehan TF, Diehl AD, Vempati UD, et al. CLO: The cell line ontology. J Biomed Semantics. 2014;5: 37.

30. Malone J, Holloway E, Adamusiak T, Kapushesky M, Zheng J, Kolesnikov N, et al. Modeling sample variables with an Experimental Factor Ontology. Bioinformatics. 2010;26: 1112–1118.

31. Mungall CJ, Torniai C, Gkoutos GV, Lewis SE, Haendel MA. Uberon, an integrative multi-species anatomy ontology. Genome Biol. 2012;13: R5.

32. Diehl AD, Meehan TF, Bradford YM, Brush MH, Dahdul WM, Dougall DS, et al. The Cell Ontology 2016: enhanced content, modularization, and ontology interoperability. J Biomed Semantics. 2016;7: 44.

33. Jeske L, Placzek S, Schomburg I, Chang A, Schomburg D. BRENDA in 2019: a European ELIXIR core data resource. Nucleic Acids Res. 2019;47: D542–D549.

34. Tonekaboni SAM, Mazrooei P, Kofia V, Haibe-Kains B, Lupien M. CREAM: Clustering of genomic REgions Analysis Method. Bioinformatics. bioRxiv; 2017. p. 958.

35. Madani Tonekaboni SA, Mazrooei P, Kofia V, Haibe-Kains B, Lupien M. Identifying clusters of cis-regulatory elements underpinning TAD structures and lineage-specific regulatory networks. Genome Res. 2019;29: 1733–1743.

36. Siepel A, Bejerano G, Pedersen JS, Hinrichs AS, Hou M, Rosenbloom K, et al. Evolutionarily conserved elements in vertebrate, insect, worm, and yeast genomes. Genome Res. 2005;15: 1034–1050.

37. Pollard KS, Hubisz MJ, Rosenbloom KR, Siepel A. Detection of nonneutral substitution rates on mammalian phylogenies. Genome Res. 2010;20: 110–121.

38. Ferré Q, Charbonnier G, Sadouni N, Lopez F, Kermezli Y, Spicuglia S, et al. OLOGRAM: Determining significance of total overlap length between genomic regions sets. Bioinformatics. 2019. doi:10.1093/bioinformatics/btz810

39. Quinlan AR, Hall IM. BEDTools: a flexible suite of utilities for comparing genomic features. Bioinformatics. 2010;26: 841–842.

40. Favorov A, Mularoni L, Cope LM, Medvedeva Y, Mironov AA, Makeev VJ, et al. Exploring massive, genome scale datasets with the GenometriCorr package. PLoS Comput Biol. 2012;8: e1002529.

41. Chen C-H, Zheng R, Tokheim C, Dong X, Fan J, Wan C, et al. Determinants of transcription factor regulatory range. Nat Commun. 2020;11: 2472.

42. ENCODE Project Consortium, Moore JE, Purcaro MJ, Pratt HE, Epstein CB, Shoresh N, et al. Expanded encyclopaedias of DNA elements in the human and mouse genomes. Nature. 2020;583: 699–710.

43. Kent WJ, Sugnet CW, Furey TS, Roskin KM, Pringle TH, Zahler AM, et al. The human genome browser at UCSC. Genome Res. 2002;12: 996–1006.

44. Andersson R, Gebhard C, Miguel-Escalada I, Hoof I, Bornholdt J, Boyd M, et al. An atlas of active enhancers across human cell types and tissues. Nature. 2014;507: 455–461.

45. Mattioli K, Volders P-J, Gerhardinger C, Lee JC, Maass PG, Melé M, et al. High-throughput functional analysis of lncRNA core promoters elucidates rules governing tissue specificity. Genome Res. 2019;29: 344–355.

46. Smith RP, Taher L, Patwardhan RP, Kim MJ, Inoue F, Shendure J, et al. Massively parallel decoding of mammalian regulatory sequences supports a flexible organizational model. Nat Genet. 2013;45: 1021–1028.

47. Andersson R, Sandelin A. Determinants of enhancer and promoter activities of regulatory elements. Nat Rev Genet. 2020;21: 71–87.

48. Theodorou V, Stark R, Menon S, Carroll JS. GATA3 acts upstream of FOXA1 in mediating ESR1 binding by shaping enhancer accessibility. Genome Res. 2013;23: 12–22.

49. Khan A, Mathelier A. Intervene: a tool for intersection and visualization of multiple gene or genomic region sets. BMC Bioinformatics. 2017;18: 287.

50. Safran M, Dalah I, Alexander J, Rosen N, Iny Stein T, Shmoish M, et al. GeneCards Version 3: the human gene integrator. Database. 2010;2010: baq020.

51. Edgar R, Domrachev M, Lash AE. Gene Expression Omnibus: NCBI gene expression and hybridization array data repository. Nucleic Acids Res. 2002;30: 207–210.

52. Raney BJ, Dreszer TR, Barber GP, Clawson H, Fujita PA, Wang T, et al. Track data hubs enable visualization of user-defined genome-wide annotations on the UCSC Genome Browser. Bioinformatics. 2014;30: 1003–1005.

53. Newman V, Moore B, Sparrow H, Perry E. The Ensembl Genome Browser: Strategies for Accessing Eukaryotic Genome Data. Methods in Molecular Biology. 2018. pp. 115–139. doi: 10.1007/978-1-4939-7737-6_6

54. Puente-Santamaria L, Wasserman WW, Del Peso L. TFEA.ChIP: a tool kit for transcription factor binding site enrichment analysis capitalizing on ChIP-seq datasets. Bioinformatics. 2019;35: 5339–5340.

55. Lachmann A, Xu H, Krishnan J, Berger SI, Mazloom AR, Ma'ayan A. ChEA: transcription factor regulation inferred from integrating genome-wide ChIP-X experiments. Bioinformatics. 2010;26: 2438–2444.

56. Verfaillie A, Imrichová H, Van de Sande B, Standaert L, Christiaens V, Hulselmans G, et al. iRegulon: from a gene list to a gene regulatory network using large motif and track collections. PLoS Comput Biol. 2014;10: e1003731.

57. Wang Z, Civelek M, Miller CL, Sheffield NC, Guertin MJ, Zang C. BART: a transcription factor prediction tool with query gene sets or epigenomic profiles. Bioinformatics. 2018;34: 2867–2869.

58. Kwon AT, Arenillas DJ, Worsley Hunt R, Wasserman WW. oPOSSUM-3: advanced analysis of regulatory motif over-representation across genes or ChIP-Seq datasets. G3. 2012;2: 987–1002.

59. Sheffield NC, Bock C. LOLA: enrichment analysis for genomic region sets and regulatory elements in R and Bioconductor. Bioinformatics. 2016;32: 587–589.

60. Fleischer T, Tekpli X, Mathelier A, Wang S, Nebdal D, Dhakal HP, et al. DNA methylation at enhancers identifies distinct breast cancer lineages. Nat Commun. 2017;8: 1379.

61. Zhao Y, Granas D, Stormo GD. Inferring binding energies from selected binding sites. PLoS Comput Biol. 2009;5: e1000590.

62. Mathelier A, Wasserman WW. The next generation of transcription factor binding site prediction. PLoS Comput Biol. 2013;9: e1003214.

63. Mathelier A, Xin B, Chiu T-P, Yang L, Rohs R, Wasserman WW. DNA Shape Features Improve Transcription Factor Binding Site Predictions In Vivo. Cell Syst. 2016;3: 278–286.e4.

64. Zhang Y, Liu T, Meyer CA, Eeckhoute J, Johnson DS, Bernstein BE, et al. Model-based analysis of ChIP-Seq (MACS). Genome Biol. 2008;9: R137.

65. Xin J, Mark A, Afrasiabi C, Tsueng G, Juchler M, Gopal N, et al. High-performance web services for querying gene and variant annotation. Genome Biol. 2016;17: 91.

66. Gupta S, Stamatoyannopoulos JA, Bailey TL, Noble WS. Quantifying similarity between motifs. Genome Biol. 2007;8: R24.

67. Bailey TL, Machanick P. Inferring direct DNA binding from ChIP-seq. Nucleic Acids Res. 2012;40: e128.

68. Pohl A, Beato M. bwtool: a tool for bigWig files. Bioinformatics. 2014;30: 1618–1619.

69. Yu G, Wang L-G, He Q-Y. ChIPseeker: an R/Bioconductor package for ChIP peak annotation, comparison and visualization. Bioinformatics. 2015;31: 2382–2383.

70. Lawrence M, Huber W, Pagès H, Aboyoun P, Carlson M, Gentleman R, et al. Software for computing and annotating genomic ranges. PLoS Comput Biol. 2013;9: e1003118.

71. Lopez F, Charbonnier G, Kermezli Y, Belhocine M, Ferré Q, Zweig N, et al. Explore, edit and leverage genomic annotations using Python GTF toolkit. Bioinformatics. 2019;35: 3487–3488.

72. Kuhn RM, Haussler D, Kent WJ. The UCSC genome browser and associated tools. Brief Bioinform. 2013;14: 144–161.

