## Supporting Material for "UniBind: maps of high-confidence direct TF-DNA interactions across nine species"

### Supplementary Tables

| Organism | Number of datasets | Number of TFs | Number of cells / tissue types | Number of TFBSs | Number of CRMs |
|---|---|---|---|---|---|
| *A. thaliana* | 121 | 41 | 27 | 181,509 | 2,226 |
| *C. elegans* | 182 | 31 | 17 | 116,018 | 604 |
| *D. rerio* | 12 | 8 | 8 | 90,455 | 1,136 |
| *D. melanogaster* | 264 | 77 | 42 | 181,359 | 1,623 |
| *H. sapiens* | 4,662 | 324 | 570 | 37,967,828 | 114,227 |
| *M. musculus* | 4,250 | 319 | 629 | 33,248,428 | 79,416 |
| *R. norvegicus* | 66 | 15 | 15 | 1,055,551 | 7,018 |
| *S. cerevisiae* | 100 | 25 | 9 | 8,152 | 207 |
| *S. pombe* | 3 | 1 | 1 | 46 | 0 |
| **Total** | 9,660 | 841 | 1,317 | 72,849,346 | 206,457 |

**Supplementary Table 1**. **Overview of the permissive collection**. Table providing the number of datasets, TFs, cell / tissue types, and TFBSs in the permissive collection of UniBind.

| Organism | Number of datasets | Number of TFs | Number of cells / tissue types | Number of TFBSs | Number of CRMs |
|---|---|---|---|---|---|
| *A. thaliana* | 78 | 33 | 22 | 169,649 | 1,262 |
| *C. elegans* | 91 | 21 | 12 | 93,138 | 503 |
| *D. rerio* | 6 | 5 | 4 | 44,187 | 617 |
| *D. melanogaster* | 109 | 22 | 28 | 95,715 | 1,162 |
| *H. sapiens* | 3,479 | 268 | 501 | 29,397,218 | 105,104 |
| *M. musculus* | 3,072 | 269 | 513 | 25,337,096 | 73,917 |
| *R. norvegicus* | 41 | 12 | 13 | 924,254 | 16,163 |
| *S. cerevisiae* | 28 | 14 | 4 | 3,776 | 117 |
| **Total** | 6,904 | 644 | 1,097 | 56,064,430 | 198,845 |

**Supplementary Table 2**. **Overview of the robust collection**. Table providing the number of datasets, TFs, cell / tissue types, and TFBSs in the robust collection of UniBind.

### Supplementary Figures

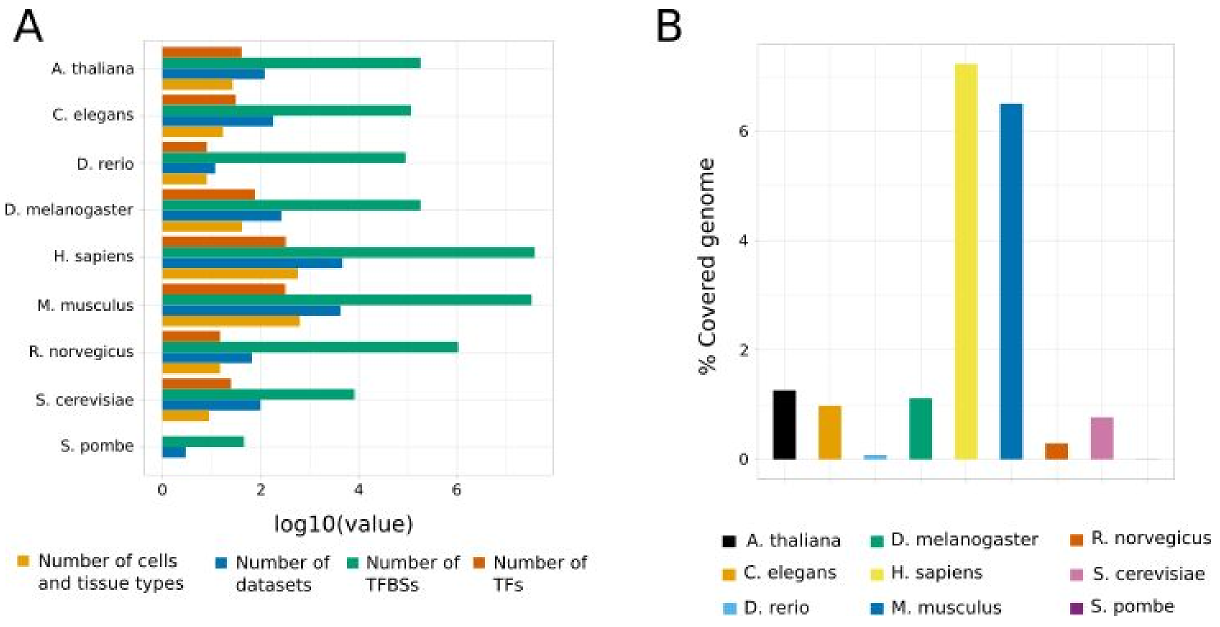


**Supplementary Figure 1**. **Visual overview of the permissive collection**. **Figure 1**. (**A**) Barplots showing the number of TFs (dark orange), TFBSs (green), datasets (blue), and cell and tissue types (light orange) stored in the permissive collection of UniBind for each analyzed species. All numbers are provided after transformation using the $log_{10}$ function. (**B**) Distribution of the percentages of the genomes covered by robust TFBSs in each species (one color per species, see legend).

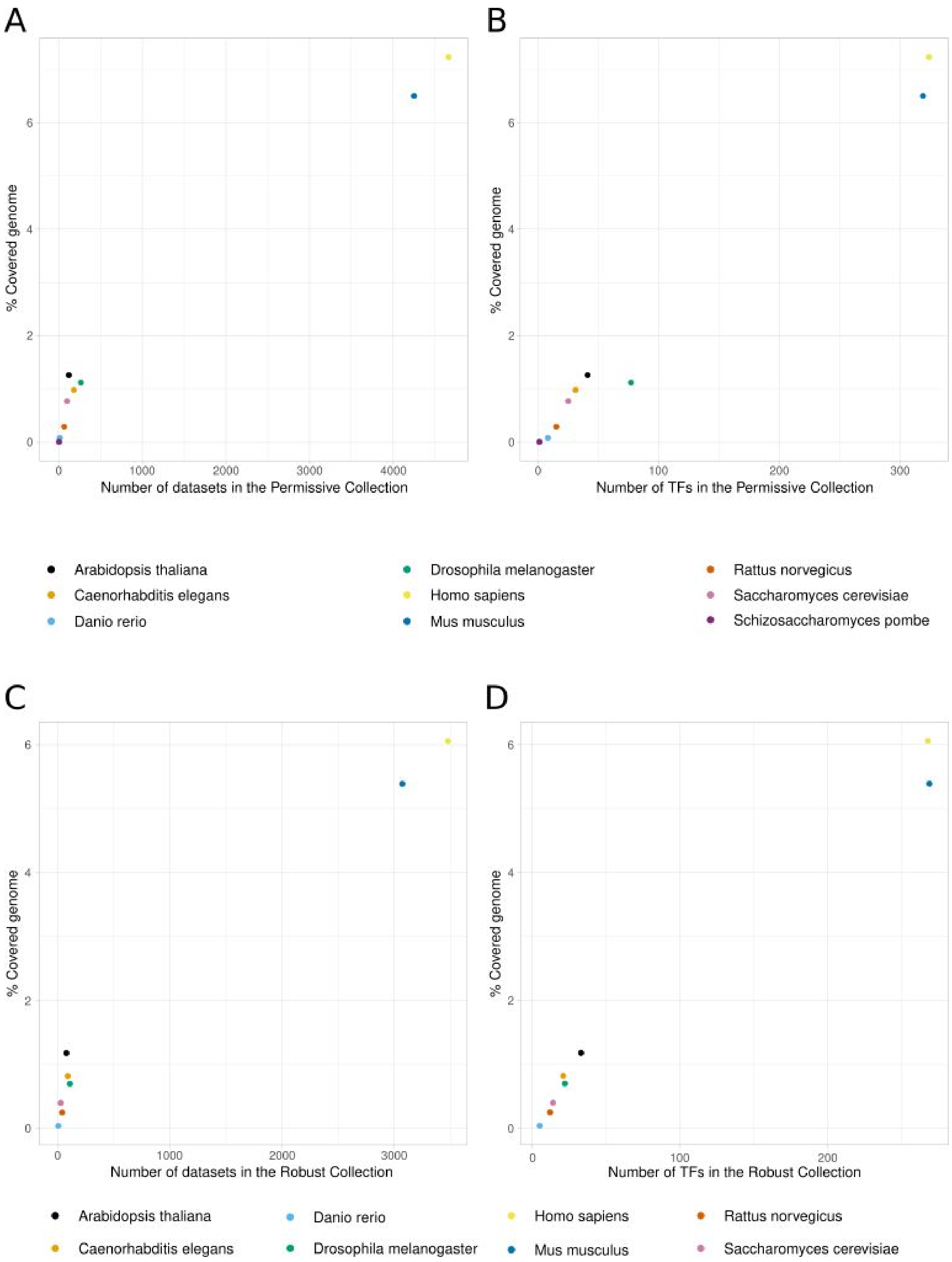


**Supplementary Figure 2**. **Relationship between number of datasets and genome coverage**. Scatter plots representing the percentage of genome coverage (y-axes) with respect to the number of datasets in the permissive (**A**) and robust (**C**) collections or the number of TFs in the permissive (**B**) and robust (**D**) collection (x-axes). Each colored point in each panel represents the data associated to one species (see legend for color coding).

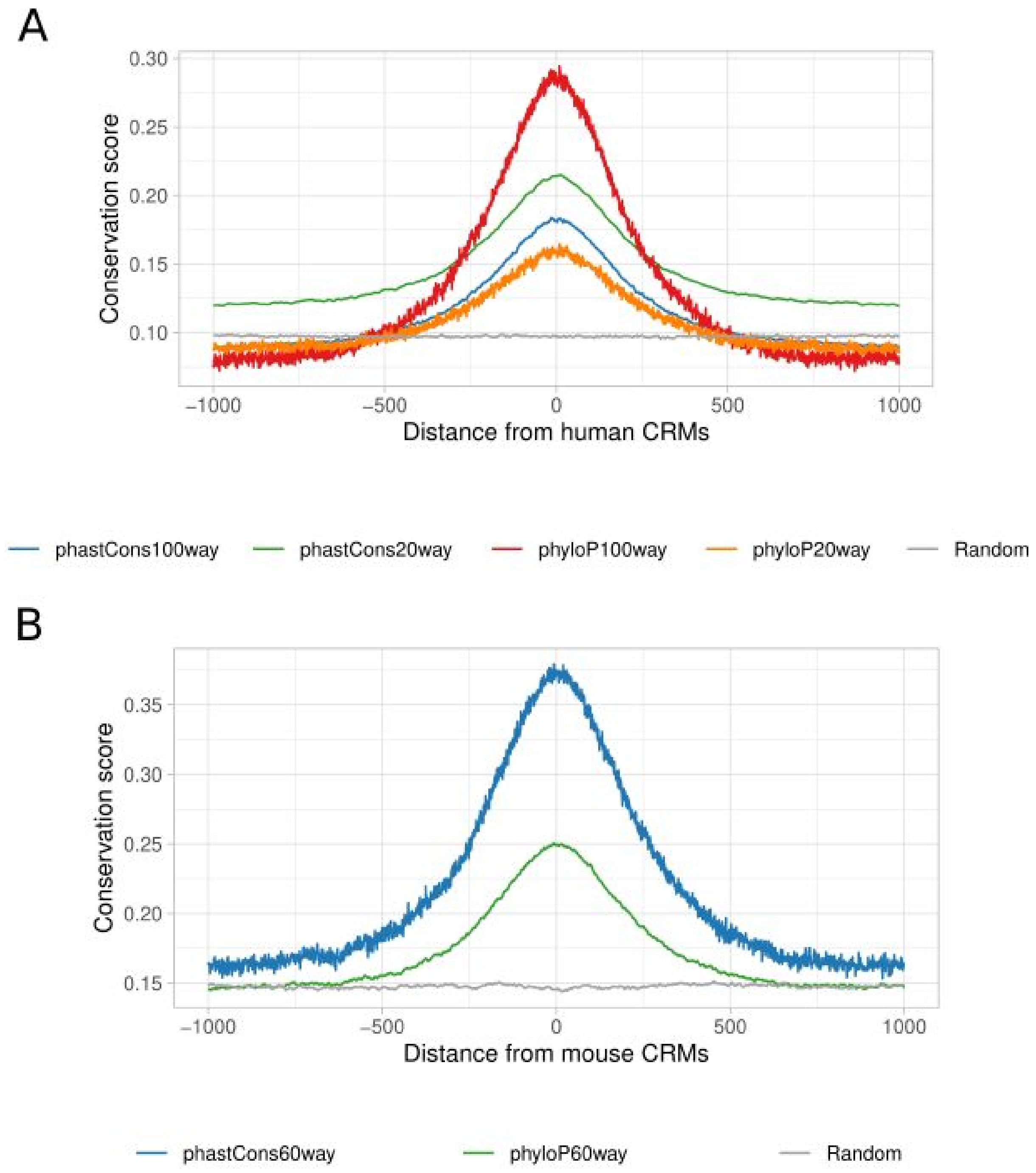


**Supplementary Figure 3**. **Evolutionary conservation at human and mouse robust CRMs**. Distributions of the average base-pair evolutionary conservation scores (phyloP and phastCons scores using multi-species genome alignments, see legend) at regions centered around UniBind human (**A**) and mouse (**B**) CRMs from the robust collection. Conservation of random CRMs was obtained by shuffling the original CRMs and obtaining the conservation score of the new regions.

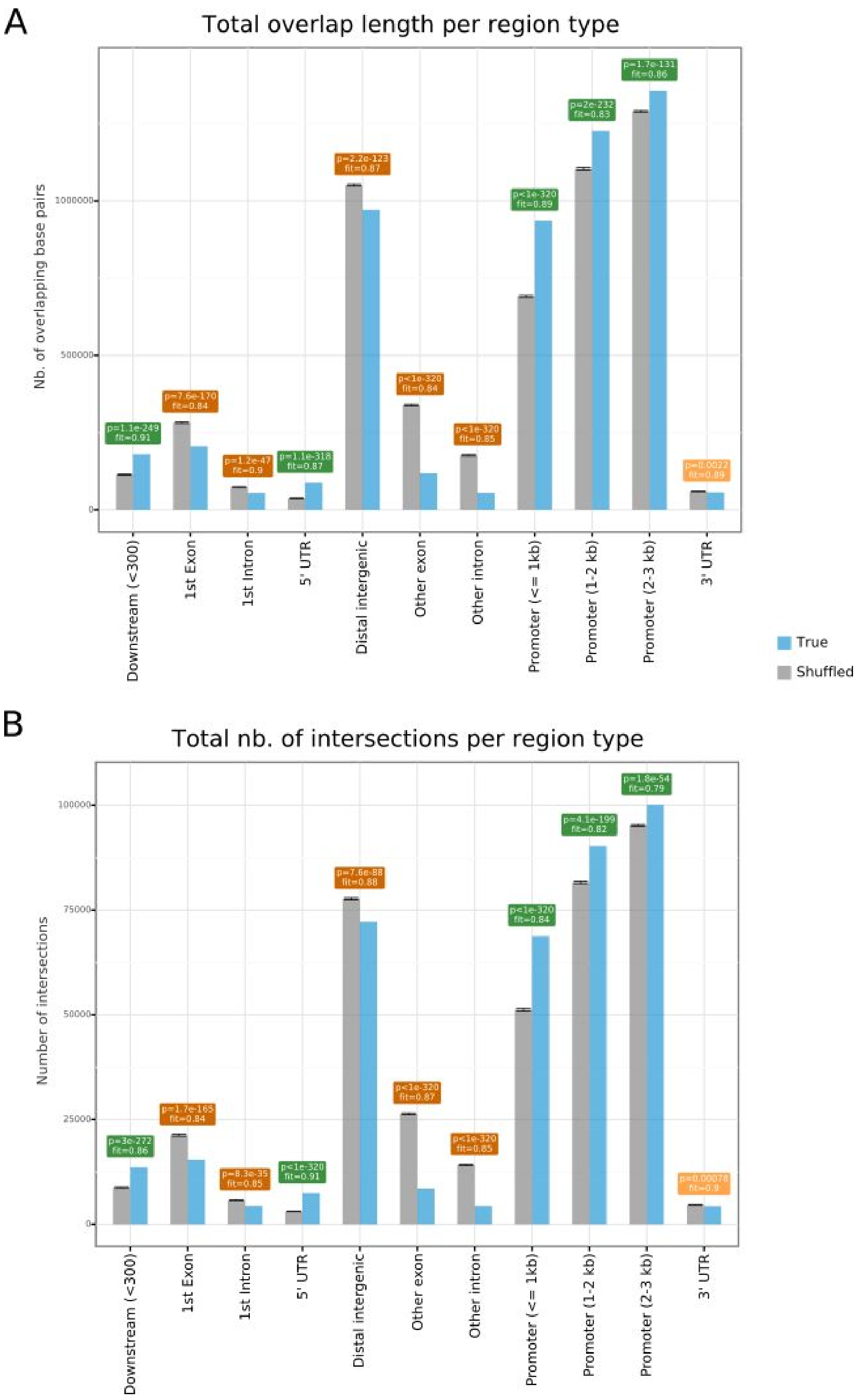


**Supplementary Figure 4**. **Enrichment analysis for *A. thaliana* TFBSs in genomic regions**. Barplots representing the expected (grey bars) versus observed (blue bars) overlap lengths (**A**) or number of intersections (**B**) between *A. thaliana* TFBSs from the robust collection and genomic annotations (x-axis). The plots and computed p-values (green: enrichment; orange: depletion) were obtained using the OLOGRAM command of the GTF toolkit.

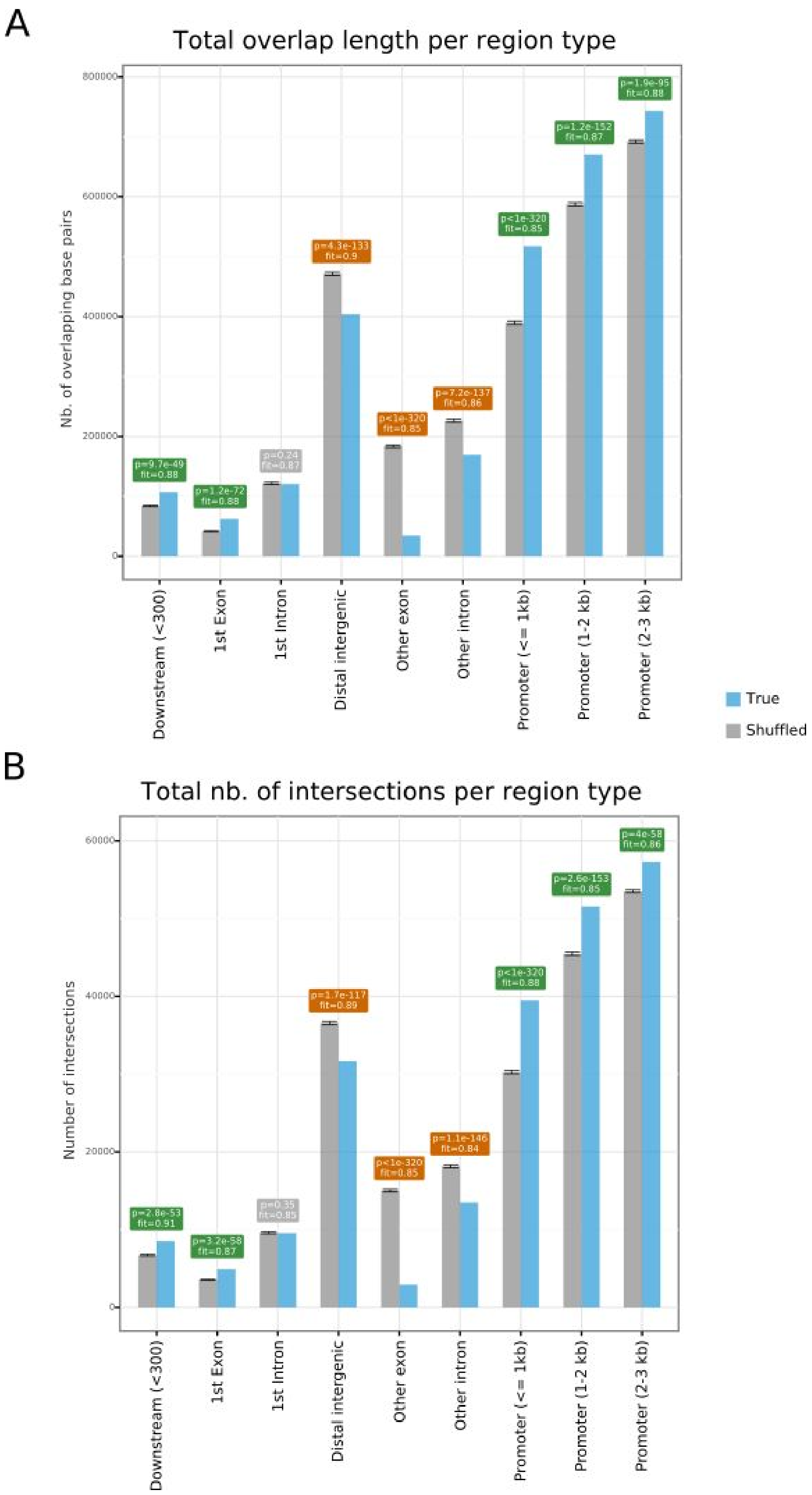


**Supplementary Figure 5**. **Enrichment analysis for *C. elegans* TFBSs in genomic regions**. Barplots representing the expected (grey bars) versus observed (blue bars) overlap lengths (**A**) or number of intersections (**B**) between *C. elegans* TFBSs from the robust collection and genomic annotations (x-axis). The plots and computed p-values (green: enrichment; orange: depletion) were obtained using the OLOGRAM command of the GTF toolkit.

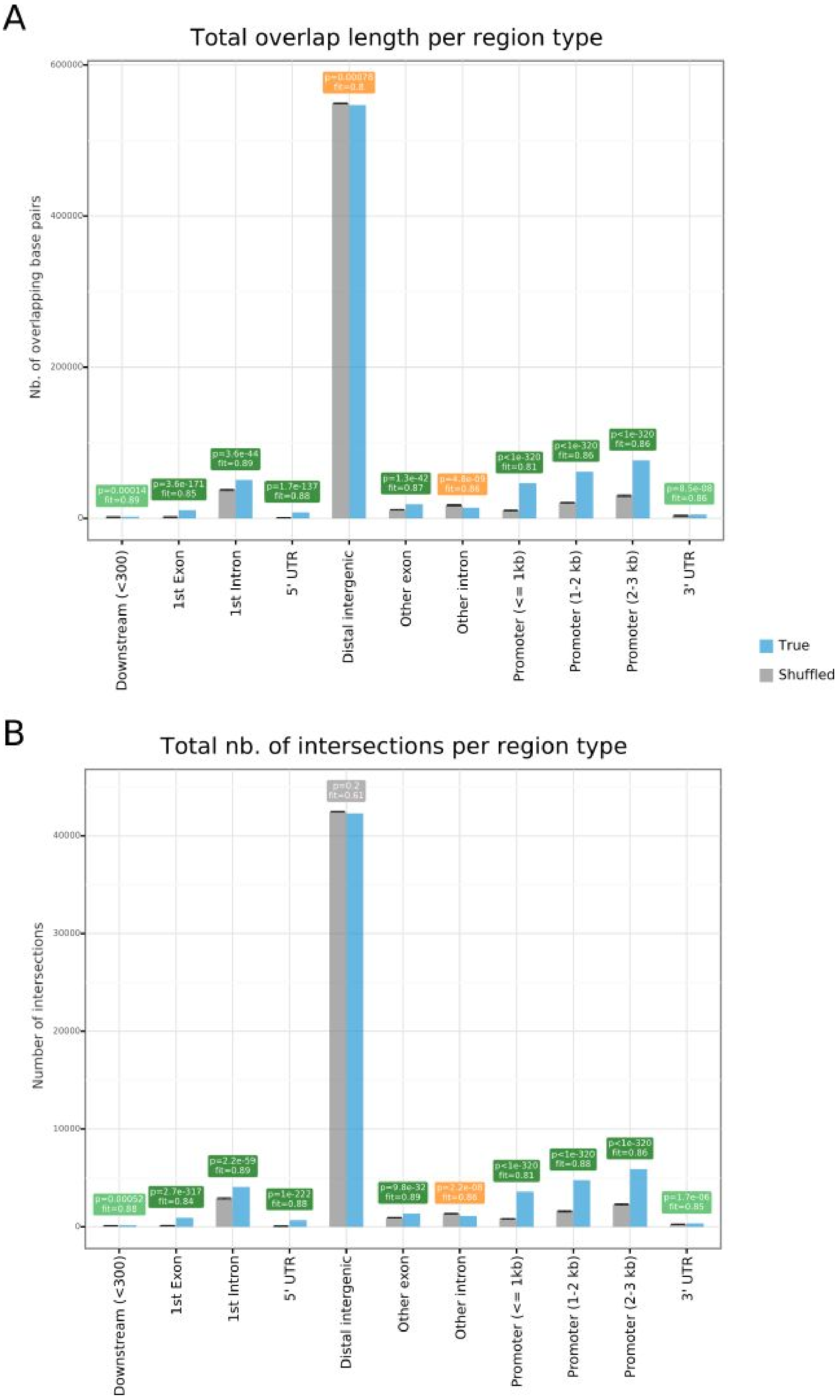


**Supplementary Figure 6**. **Enrichment analysis for *D. rerio* TFBSs in genomic regions**. Barplots representing the expected (grey bars) versus observed (blue bars) overlap lengths (**A**) or number of intersections (**B**) between *D. rerio* TFBSs from the robust collection and genomic annotations (x-axis). The plots and computed p-values (green: enrichment; orange: depletion) were obtained using the OLOGRAM command of the GTF toolkit.

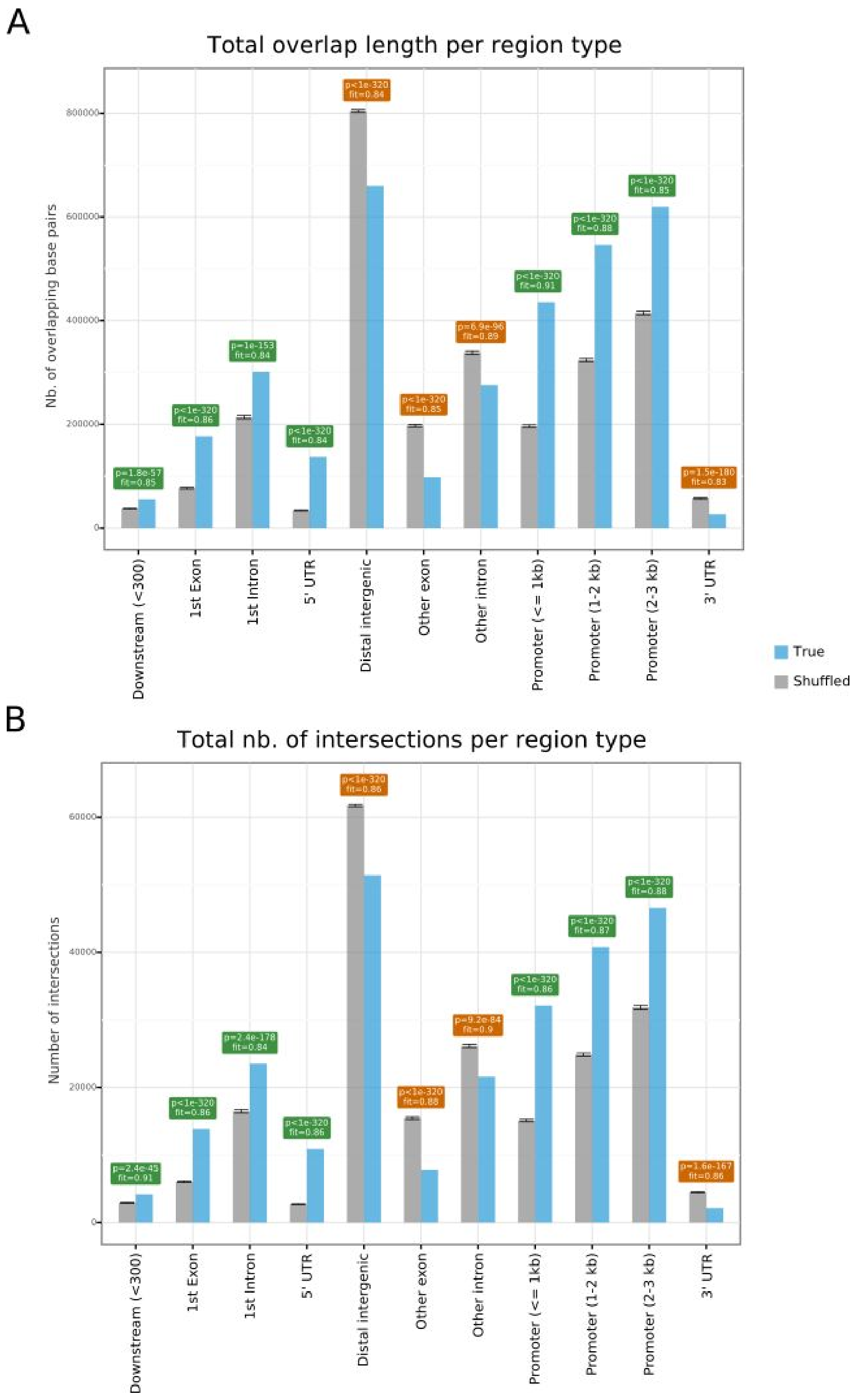


**Supplementary Figure 7**. **Enrichment analysis for *D. melanogaster* TFBSs in genomic regions**. Barplots representing the expected (grey bars) versus observed (blue bars) overlap lengths (**A**) or number of intersections (**B**) between *D. melanogaster* TFBSs from the robust collection and genomic annotations (x-axis). The plots and computed p-values (green: enrichment; orange: depletion) were obtained using the OLOGRAM command of the GTF toolkit.

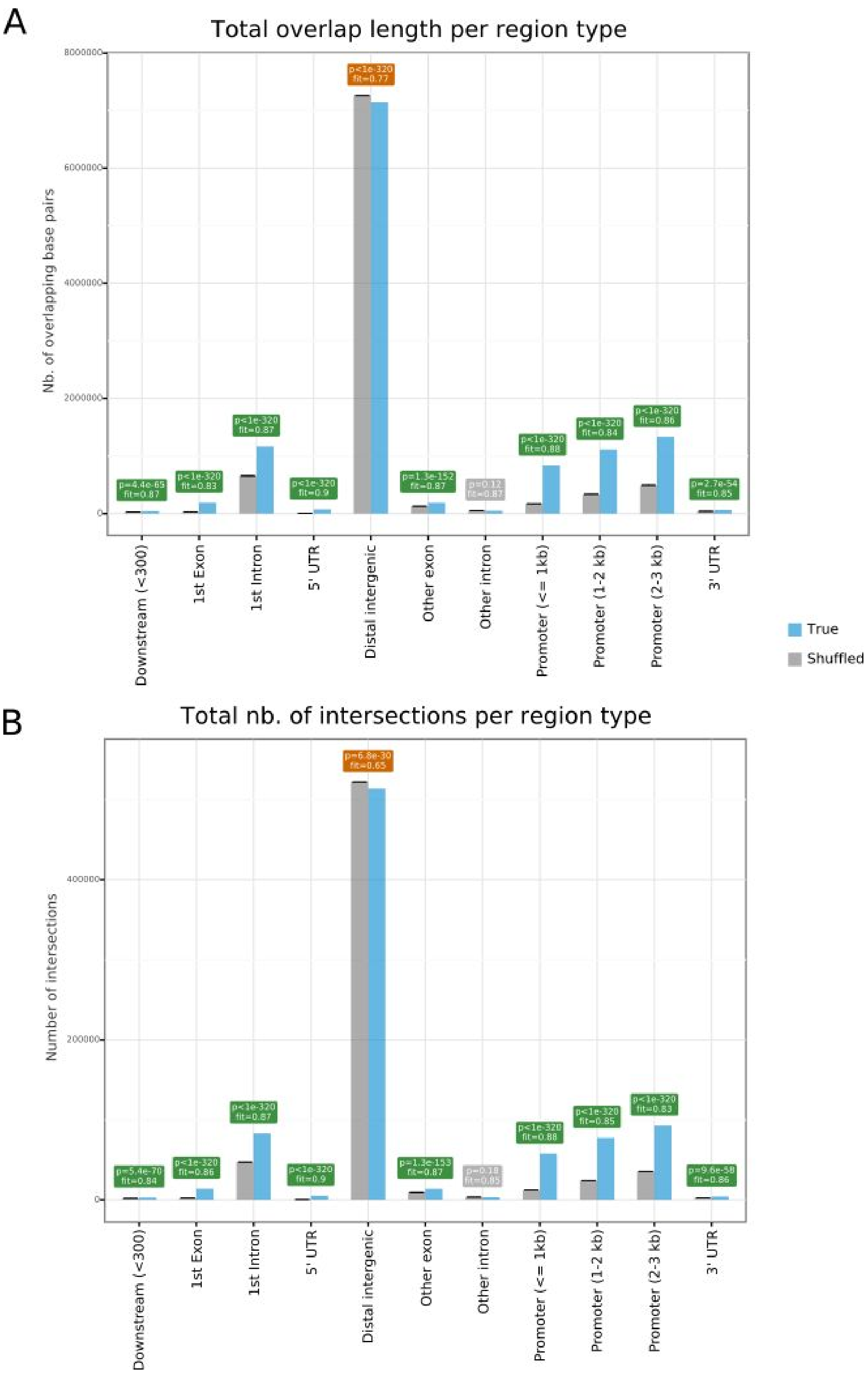


**Supplementary Figure 8**. **Enrichment analysis for *R. norvegicus* TFBSs in genomic regions**. Barplots representing the expected (grey bars) versus observed (blue bars) overlap lengths (**A**) or number of intersections (**B**) between *R. norvegicus* TFBSs from the robust collection and genomic annotations (x-axis). The plots and computed p-values (green: enrichment; orange: depletion) were obtained using the OLOGRAM command of the GTF toolkit.

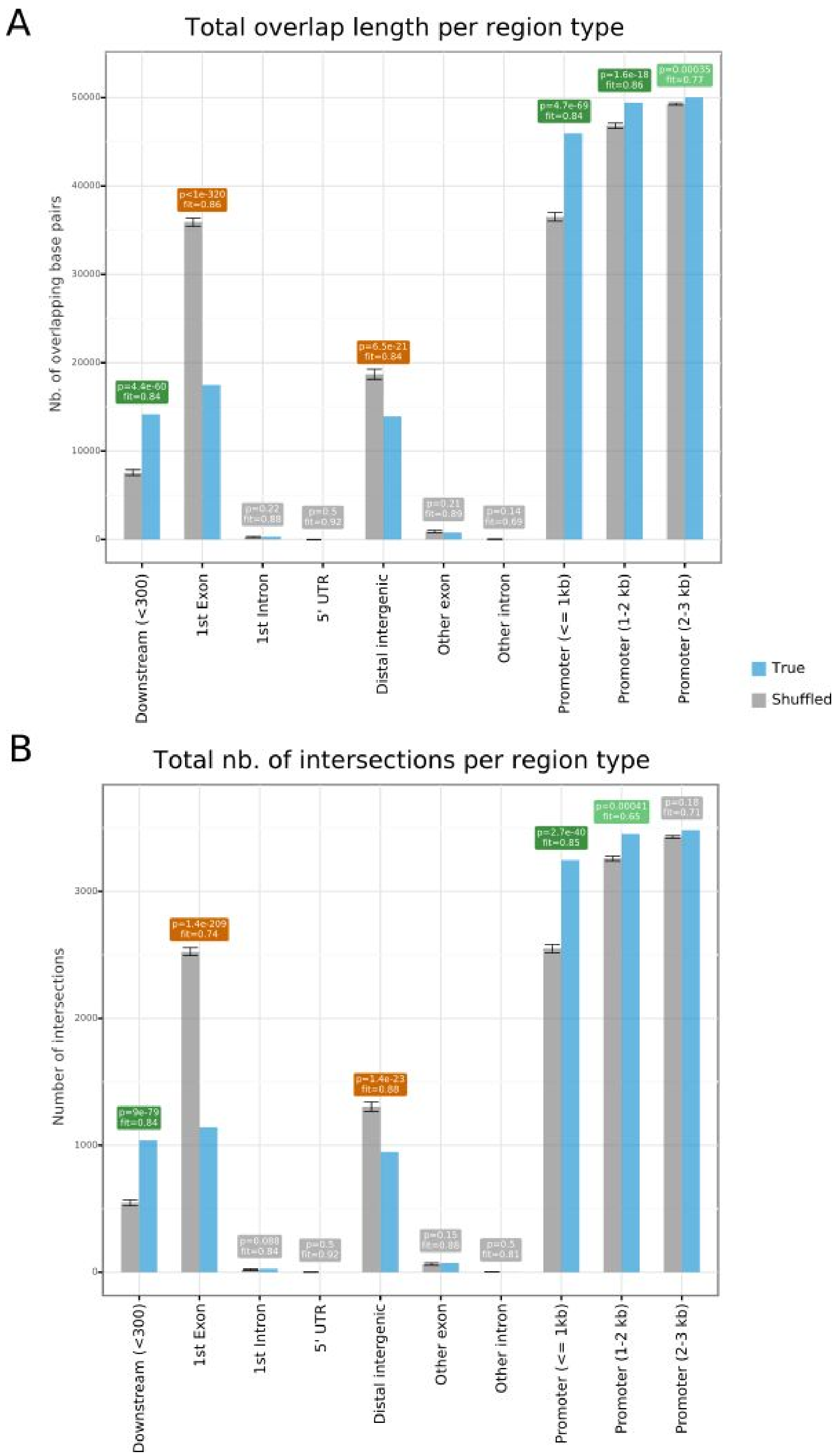


**Supplementary Figure 9**. **Enrichment analysis for *S. cerevisae* TFBSs in genomic regions**. Barplots representing the expected (grey bars) versus observed (blue bars) overlap lengths (**A**) or number of intersections (**B**) between *S. cerevisae* TFBSs from the robust collection and genomic annotations (x-axis). The plots and computed p-values (green: enrichment; orange: depletion) were obtained using the OLOGRAM command of the GTF toolkit.

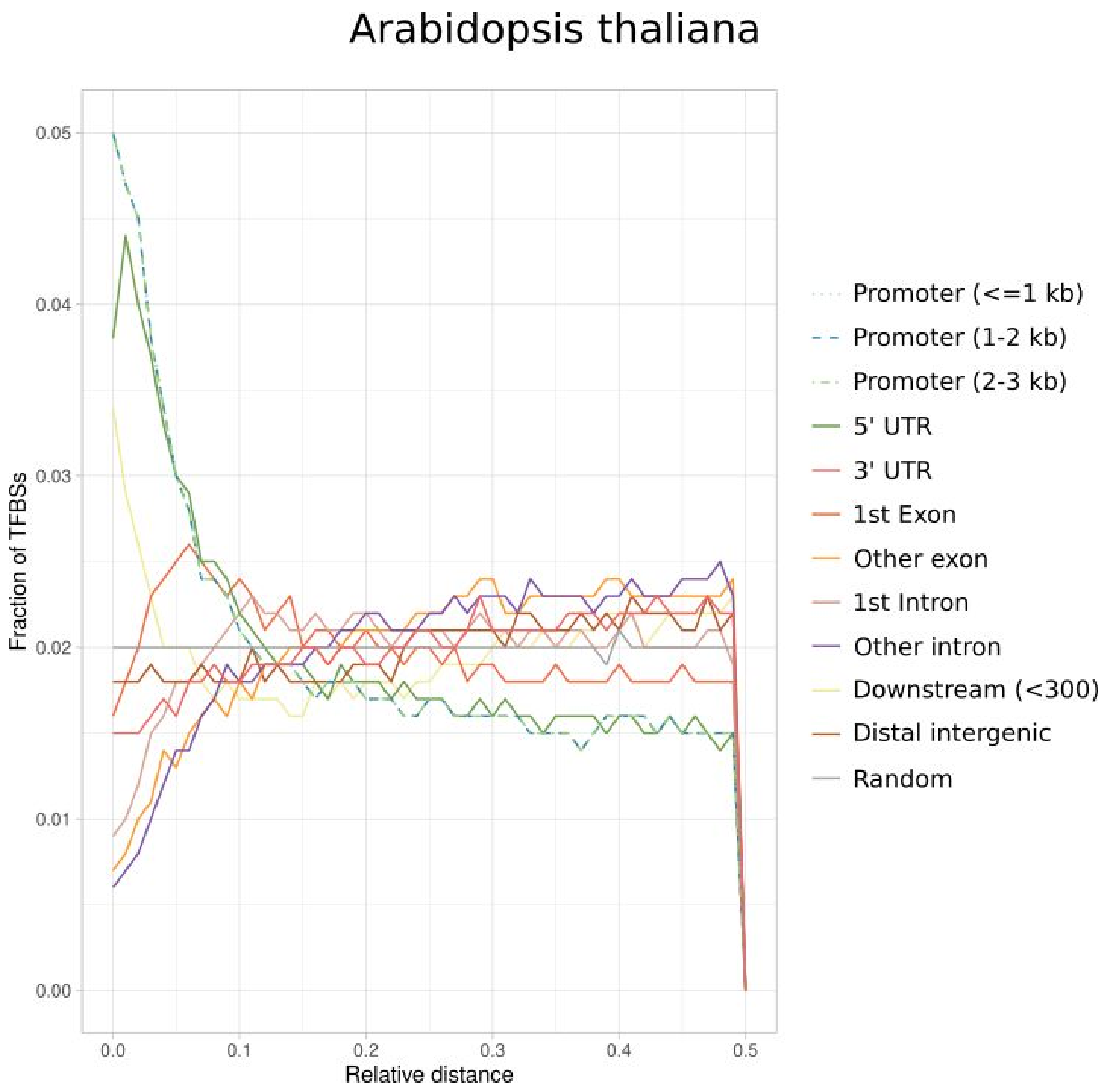


**Supplementary Figure 10**. **Analysis of the overlap of TFBSs with respect to genomic annotations in *Arabidopsis thaliana***. Fraction of TFBSs in the UniBind robust collection (y-axis) with respect to increasing relative distances (x-axis) from different genomic regions computed using the *bedtools reldist* command. When two genomic tracks are not spatially related, one expects the fraction of relative distance distribution to be uniform.

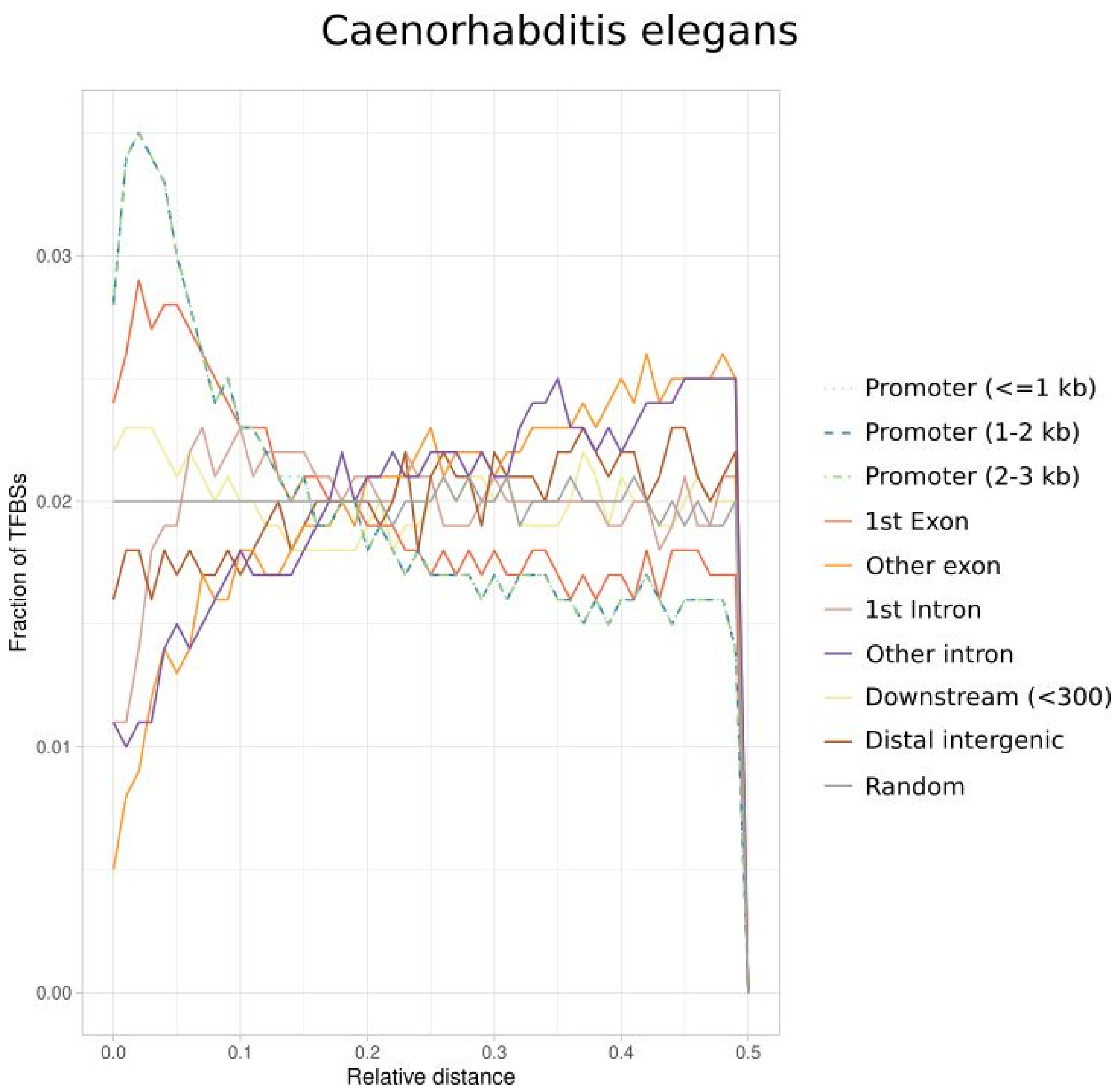


**Supplementary Figure 11**. **Analysis of the overlap of TFBSs with respect to genomic annotations in *Caenorhabditis elegans***. Fraction of TFBSs in the UniBind robust collection (y-axis) with respect to increasing relative distances (x-axis) from different genomic regions computed using the *bedtools reldist* command. When two genomic tracks are not spatially related, one expects the fraction of relative distance distribution to be uniform.

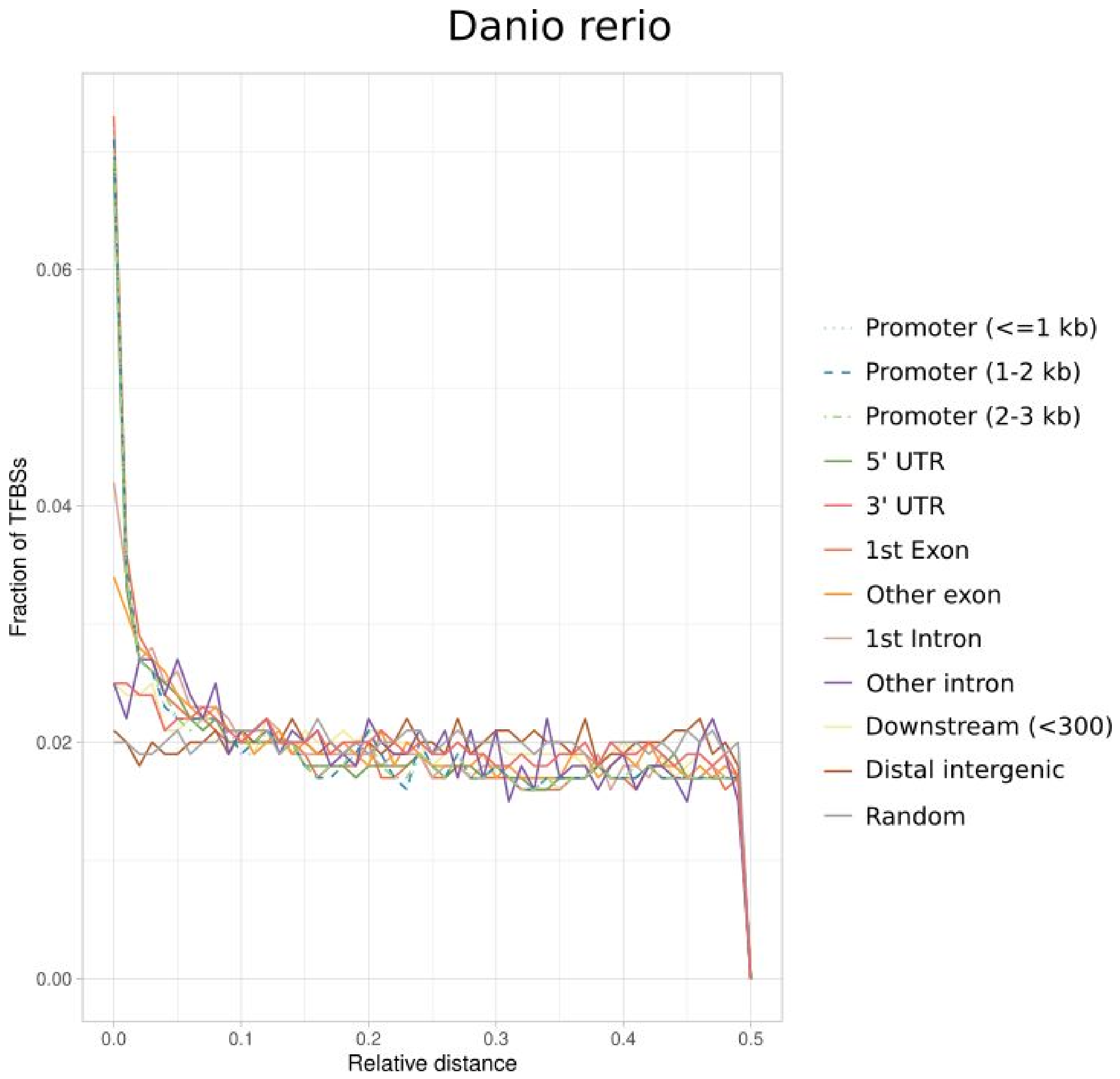


**Supplementary Figure 12**. **Analysis of the overlap of TFBSs with respect to genomic annotations in *Danio rerio***. Fraction of TFBSs in the UniBind robust collection (y-axis) with respect to increasing relative distances (x-axis) from different genomic regions computed using the *bedtools reldist* command. When two genomic tracks are not spatially related, one expects the fraction of relative distance distribution to be uniform.

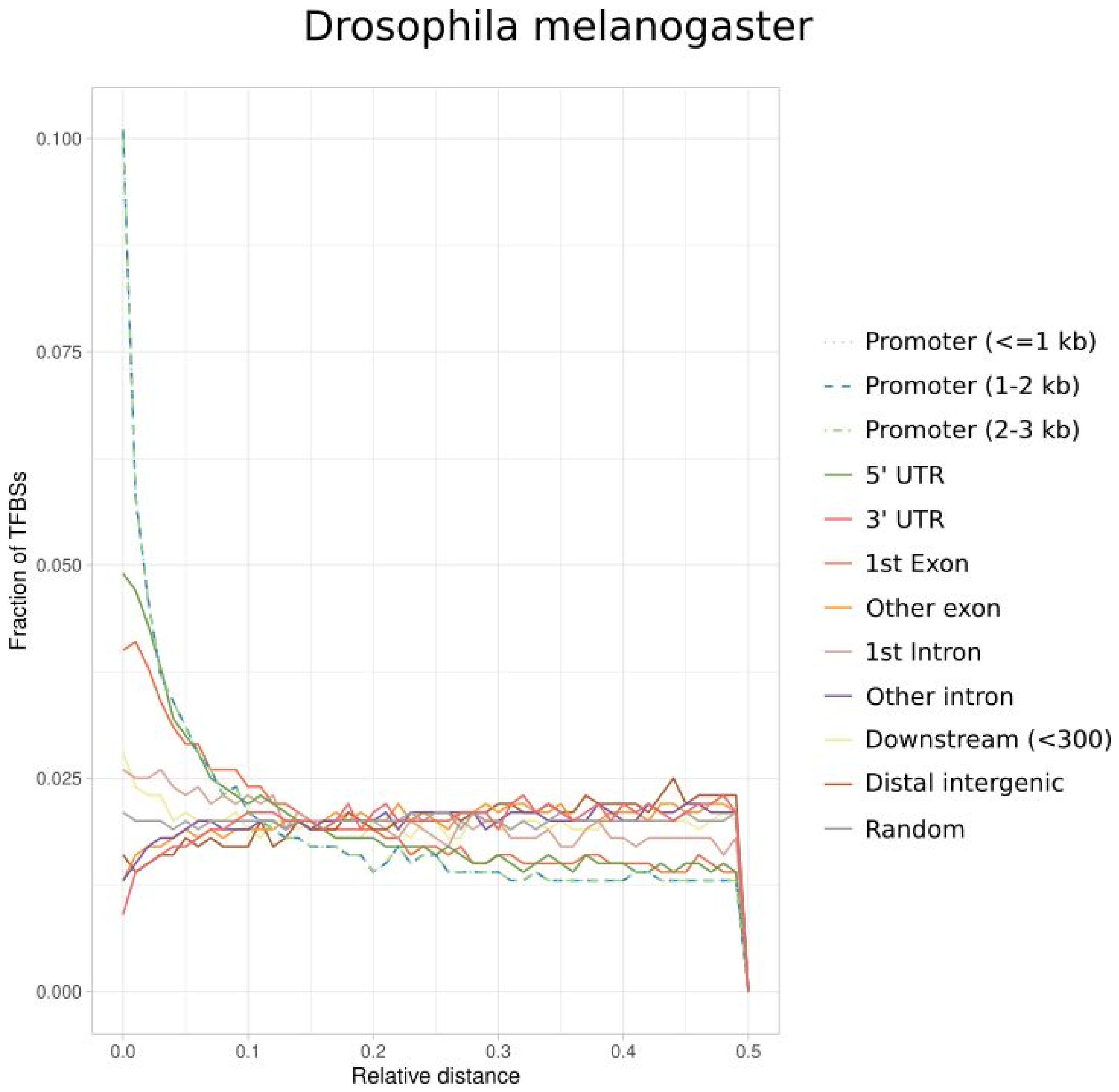


**Supplementary Figure 13**. **Analysis of the overlap of TFBSs with respect to genomic annotations in *Drosophila melanogaster***. Fraction of TFBSs in the UniBind robust collection (y-axis) with respect to increasing relative distances (x-axis) from different genomic regions computed using the *bedtools reldist* command. When two genomic tracks are not spatially related, one expects the fraction of relative distance distribution to be uniform.

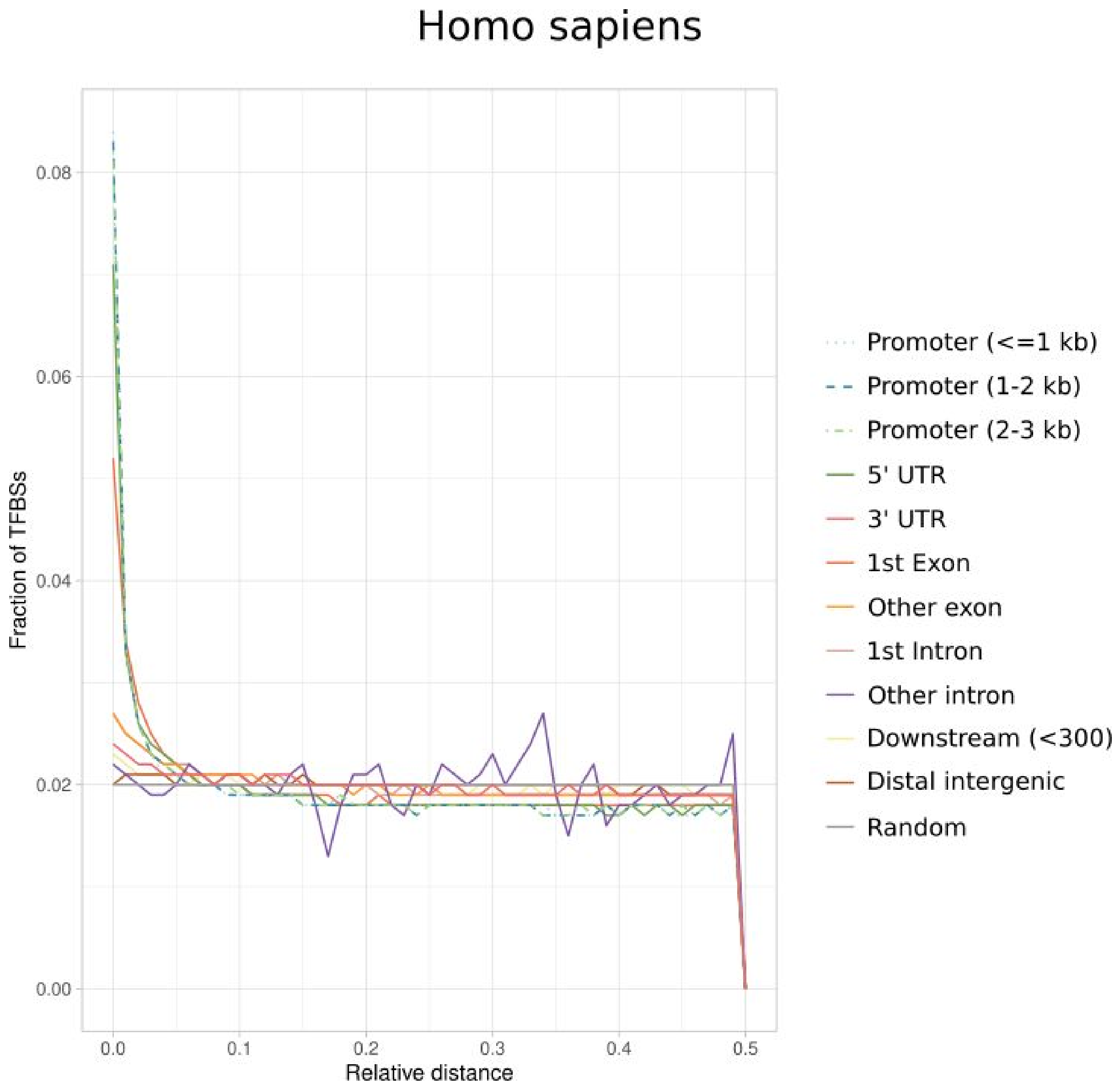


**Supplementary Figure 14**. **Analysis of the overlap of TFBSs with respect to genomic annotations in *Homo sapiens***. Fraction of TFBSs in the UniBind robust collection (y-axis) with respect to increasing relative distances (x-axis) from different genomic regions computed using the *bedtools reldist* command. When two genomic tracks are not spatially related, one expects the fraction of relative distance distribution to be uniform.

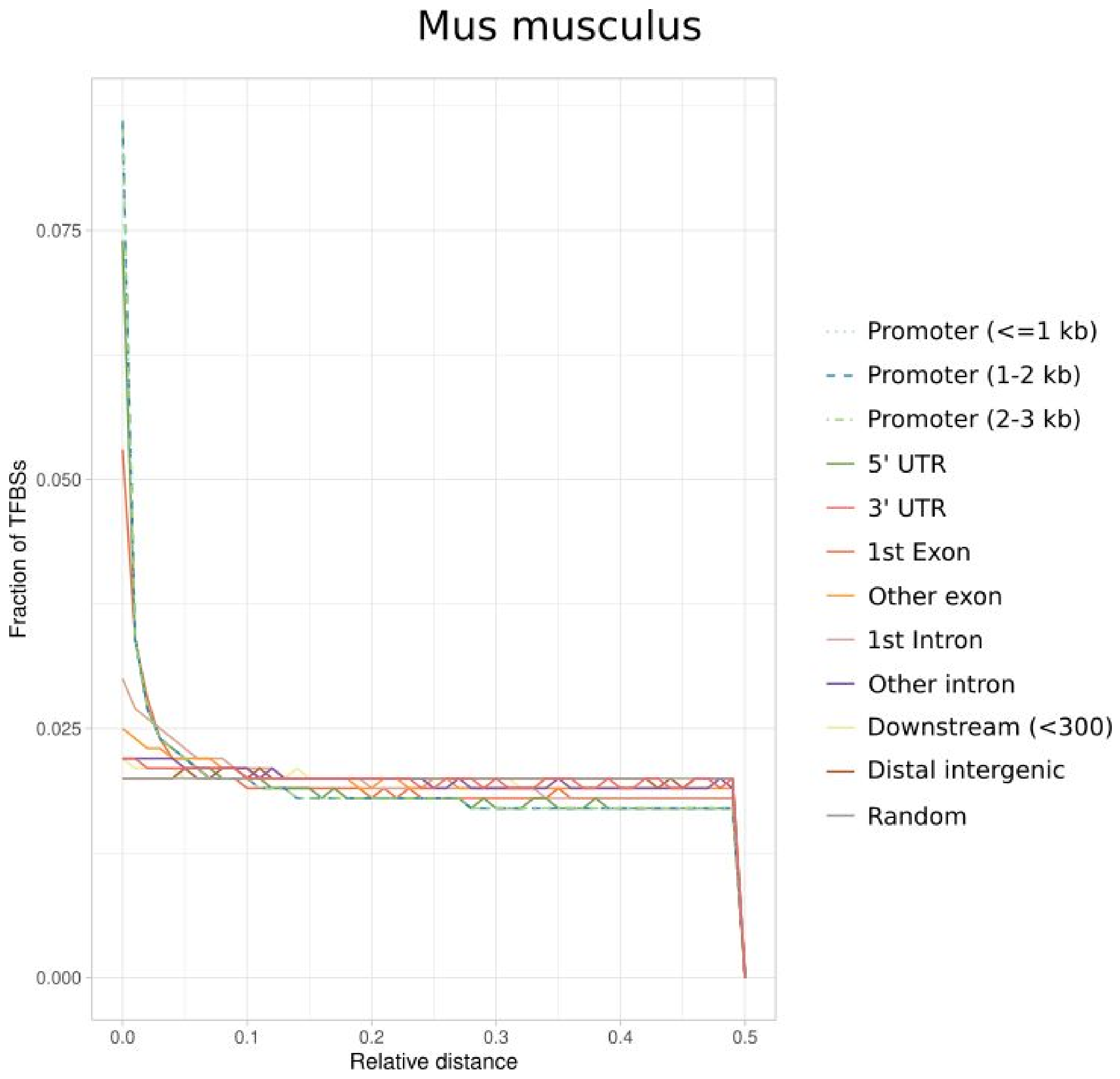


**Supplementary Figure 15**. **Analysis of the overlap of TFBSs with respect to genomic annotations in *Mus musculus***. Fraction of TFBSs in the UniBind robust collection (y-axis) with respect to increasing relative distances (x-axis) from different genomic regions computed using the *bedtools reldist* command. When two genomic tracks are not spatially related, one expects the fraction of relative distance distribution to be uniform.

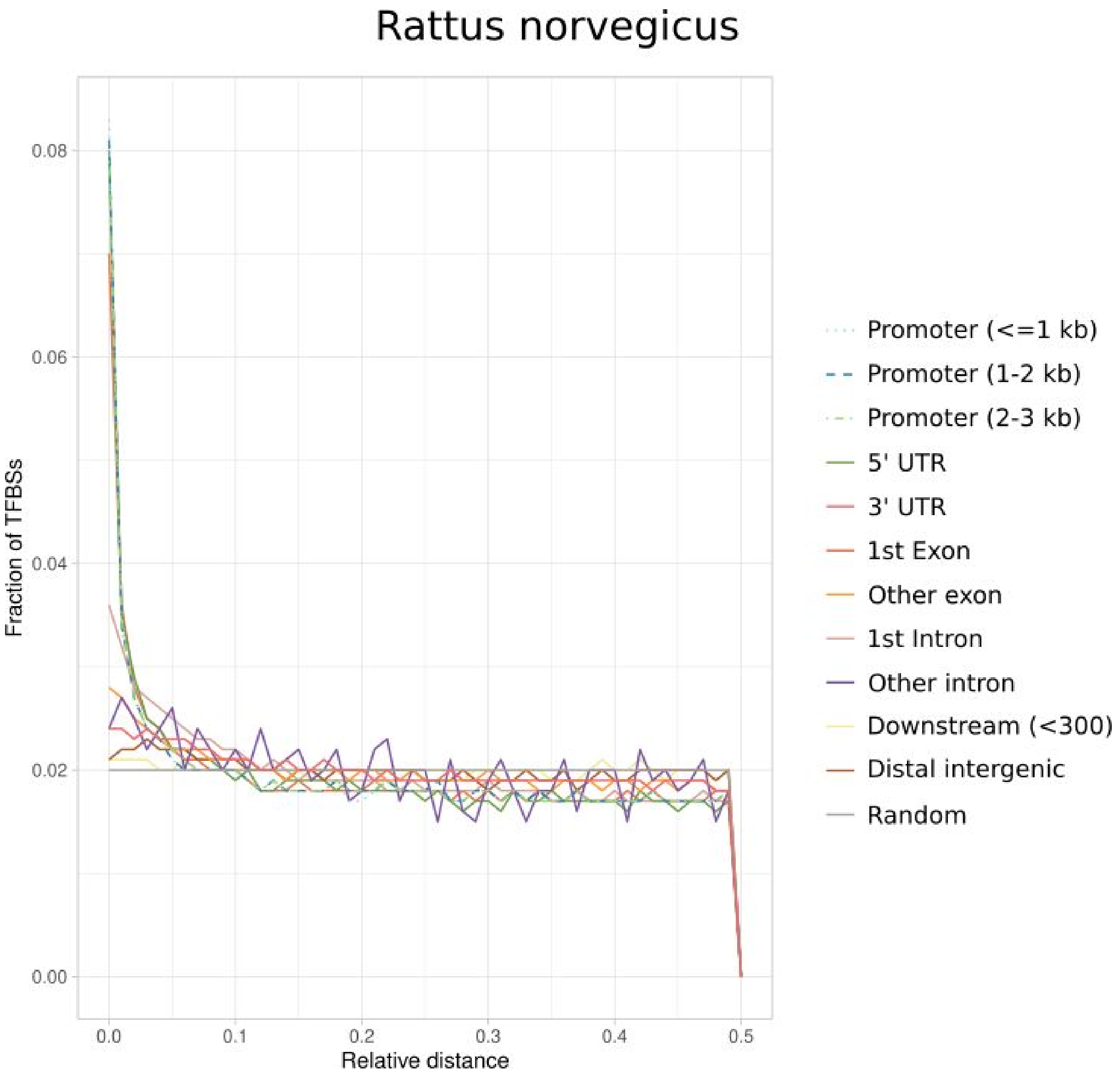


**Supplementary Figure 16**. **Analysis of the overlap of TFBSs with respect to genomic annotations in *Rattus norvegicus***. Fraction of TFBSs in the UniBind robust collection (y-axis) with respect to increasing relative distances (x-axis) from different genomic regions computed using the *bedtools reldist* command. When two genomic tracks are not spatially related, one expects the fraction of relative distance distribution to be uniform.

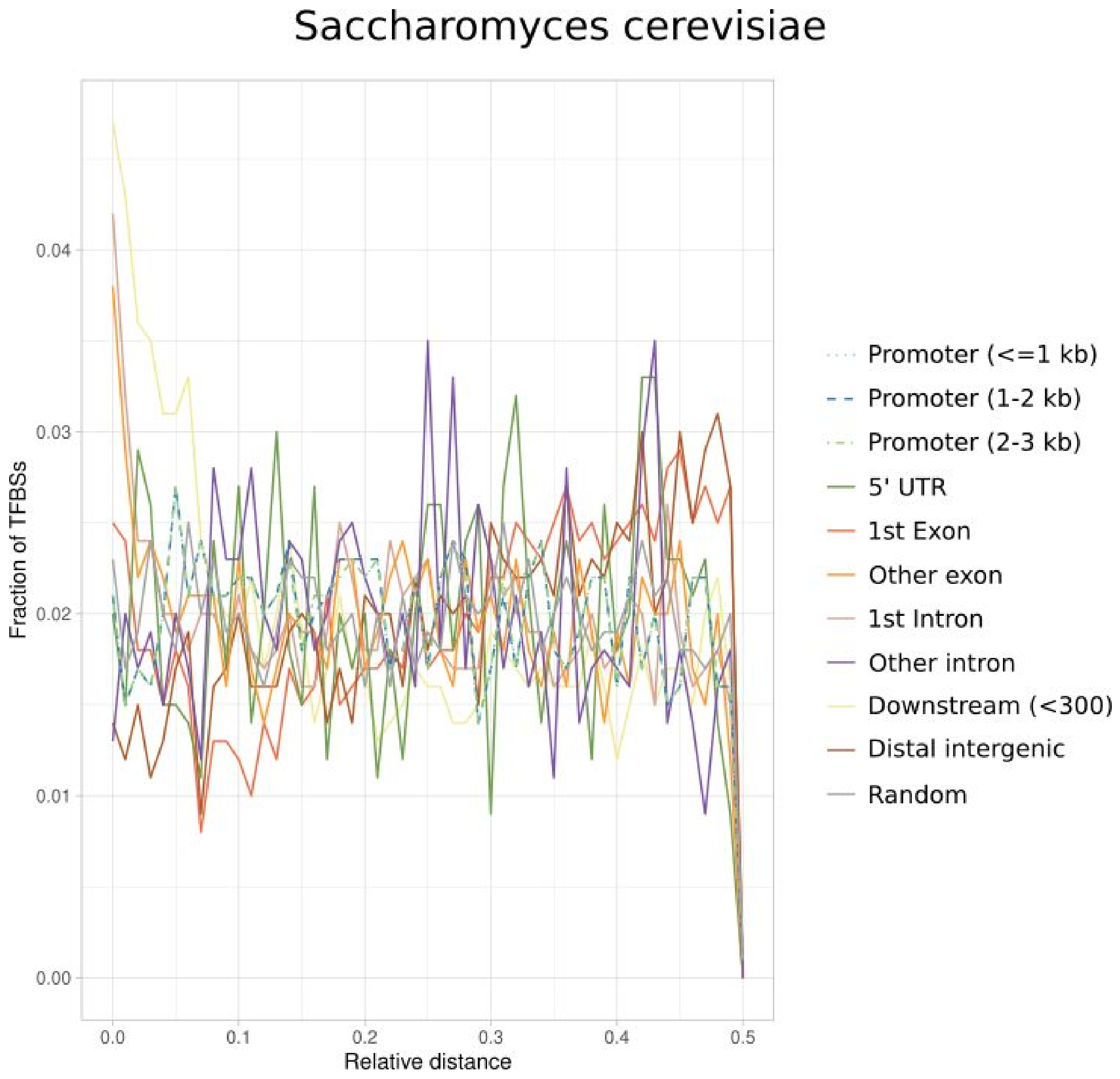


**Supplementary Figure 17**. **Analysis of the overlap of TFBSs with respect to genomic annotations in *Saccharomyces cerevisiae***. Fraction of TFBSs in the UniBind robust collection (y-axis) with respect to increasing relative distances (x-axis) from different genomic regions computed using the *bedtools reldist* command. When two genomic tracks are not spatially related, one expects the fraction of relative distance distribution to be uniform.

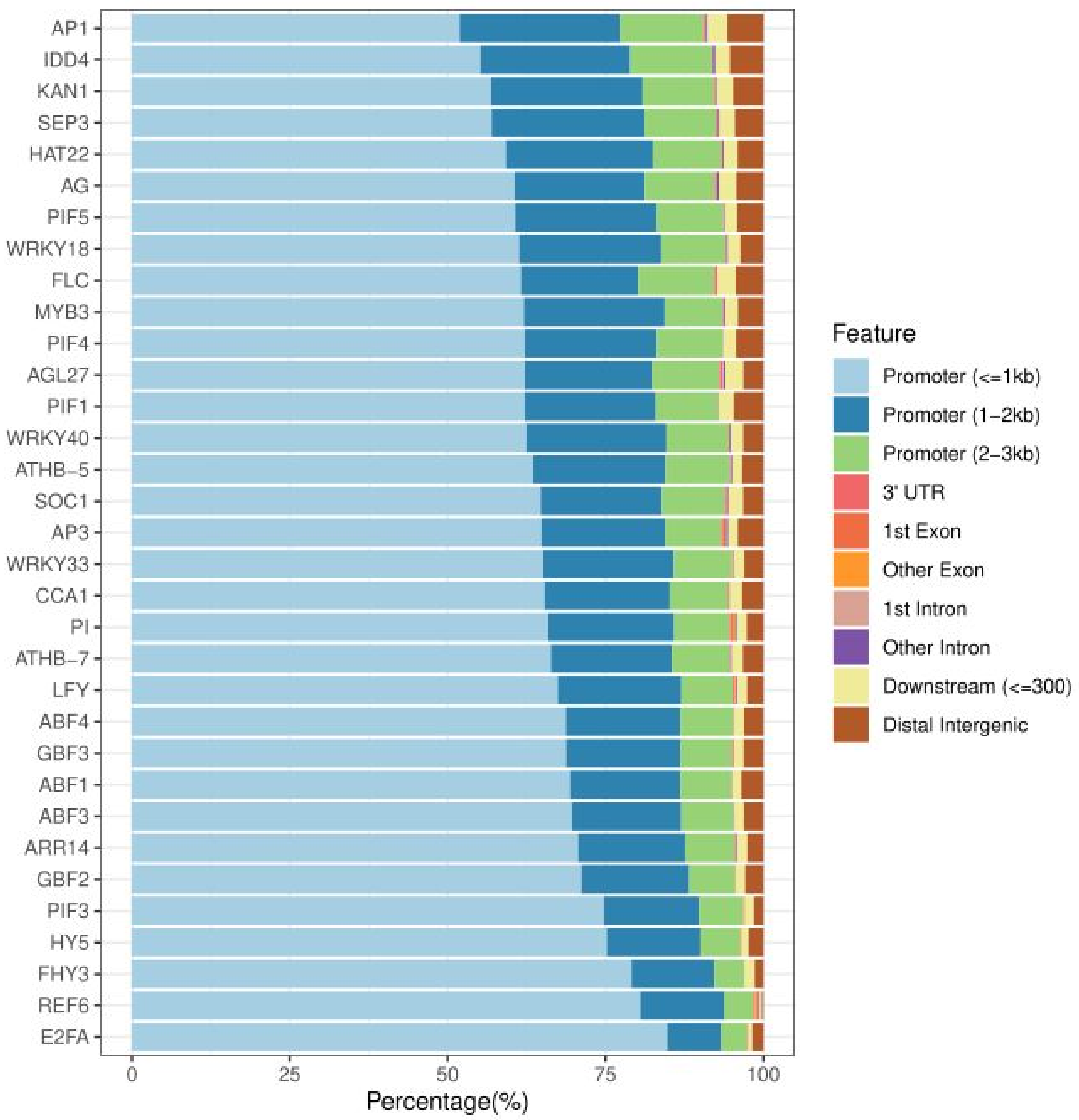


**Supplementary Figure 18. Genomic distribution of *A. thaliana* TFBSs**. Distribution of the proportion of *A. thaliana* UniBind robust TFBSs overlapping with different types of genomic regions (columns; see legend) across TFs (rows).

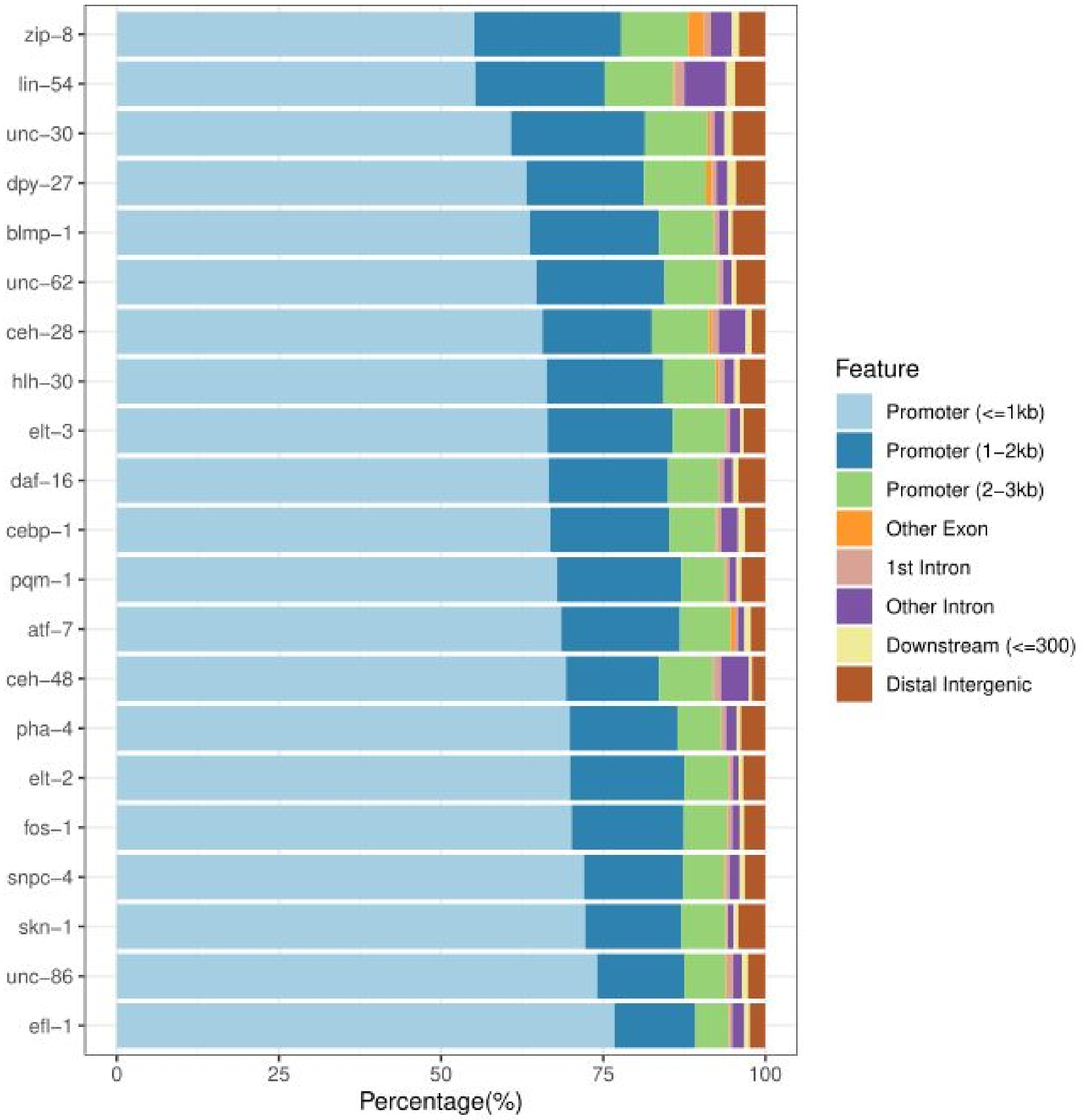


**Supplementary Figure 19. Genomic distribution of *C. elegans* TFBSs**. Distribution of the proportion of *C. elegans* UniBind robust TFBSs overlapping with different types of genomic regions (columns; see legend) across TFs (rows).

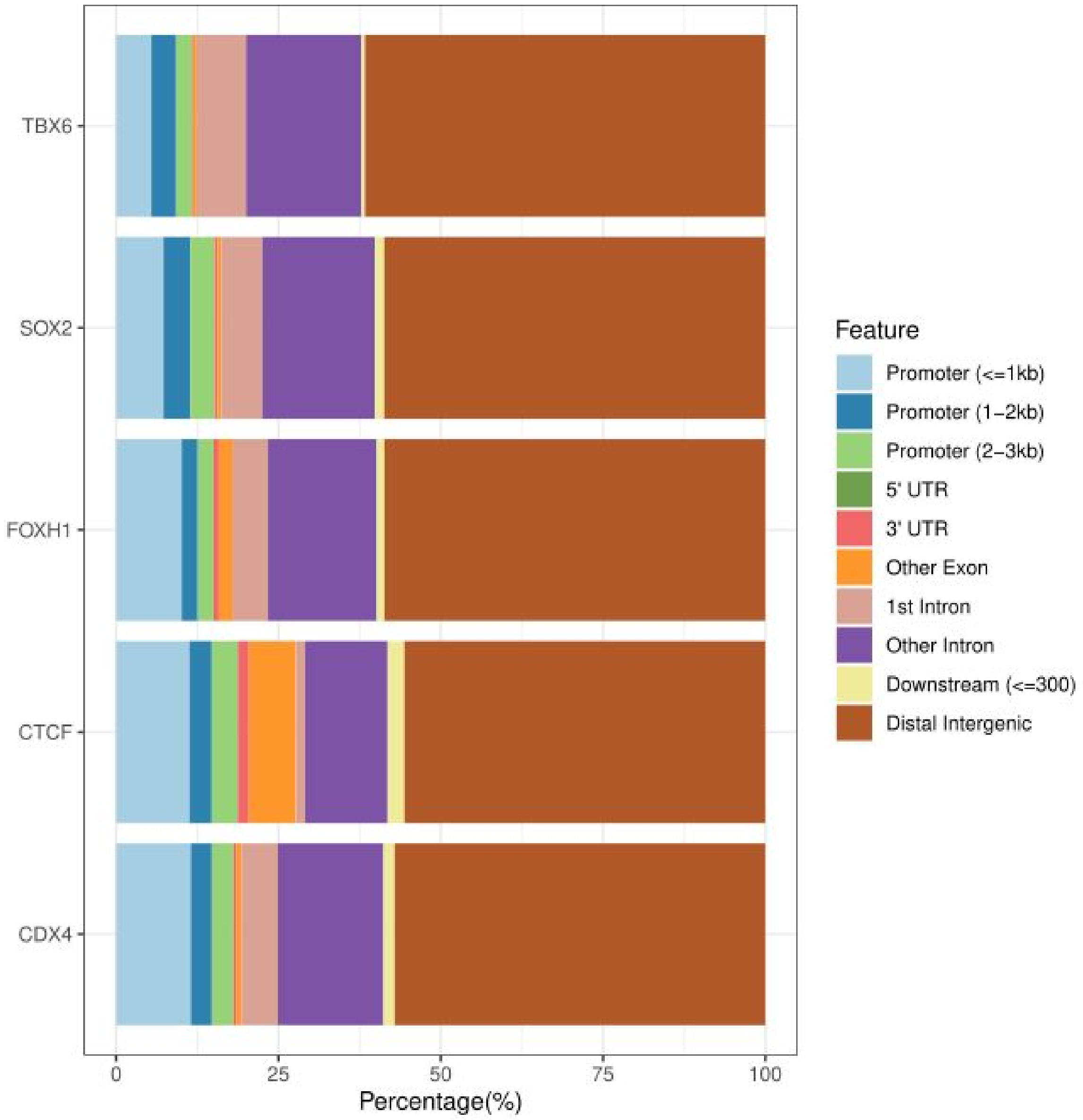


**Supplementary Figure 20. Genomic distribution of *D. rerio* TFBSs**. Distribution of the proportion of *D. rerio* UniBind robust TFBSs overlapping with different types of genomic regions (columns; see legend) across TFs (rows).

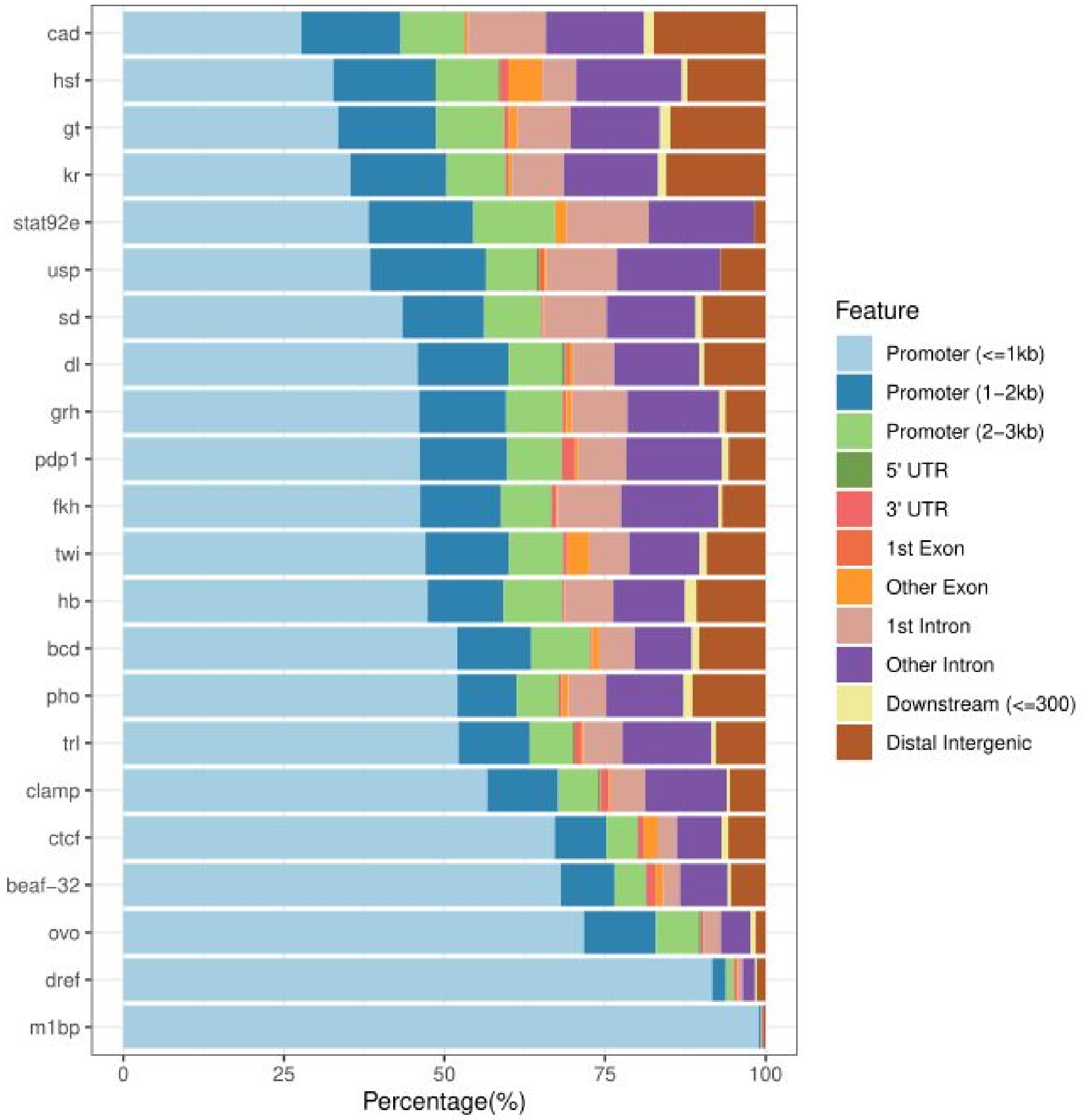


**Supplementary Figure 21. Genomic distribution of *D. melanogaster* TFBSs**. Distribution of the proportion of *D. melanogaster* UniBind robust TFBSs overlapping with different types of genomic regions (columns; see legend) across TFs (rows).

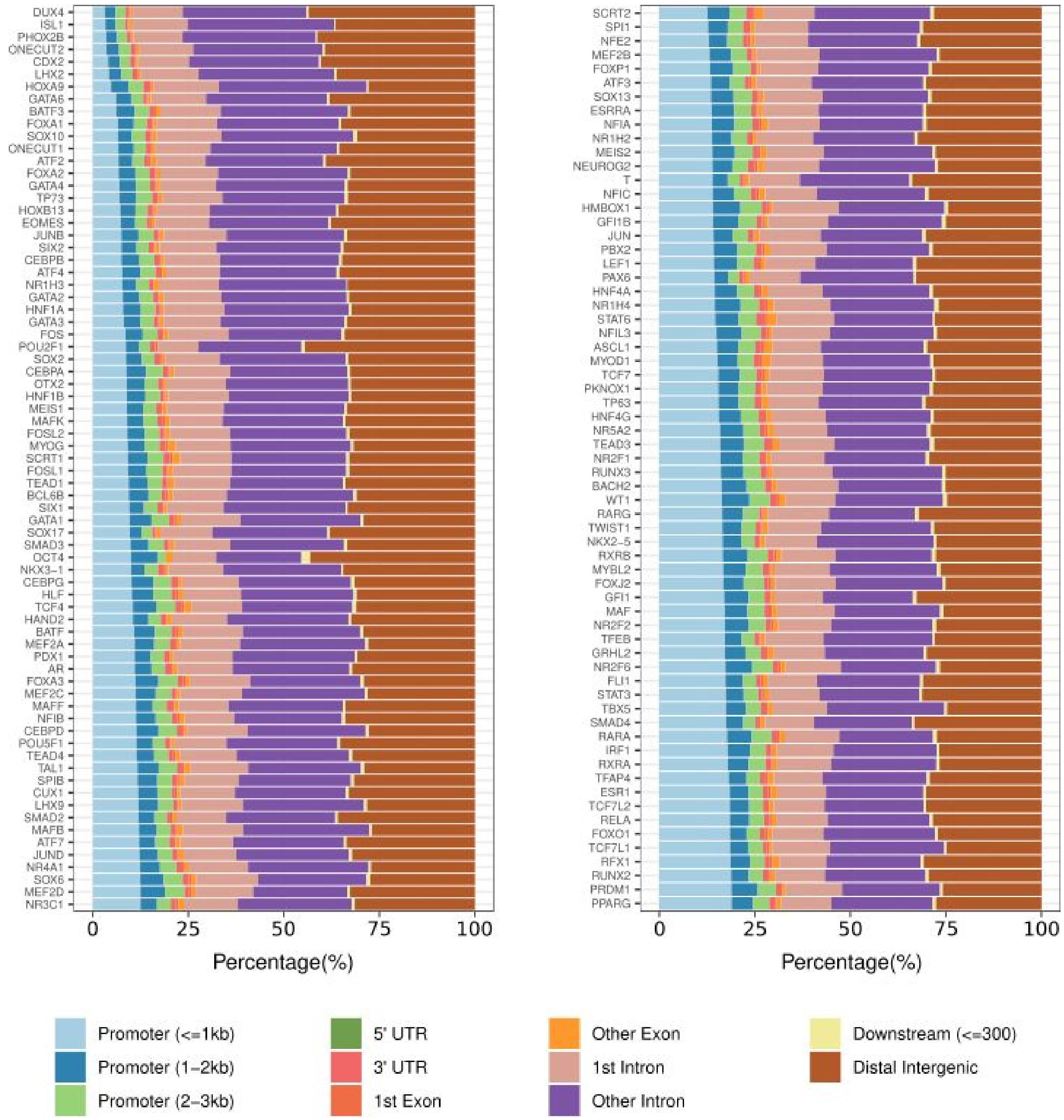


**Supplementary Figure 22. Genomic distribution of *H. sapiens* TFBSs**. Distribution of the proportion of *H. sapiens* UniBind robust TFBSs overlapping with different types of genomic regions (columns; see legend) across TFs (rows).

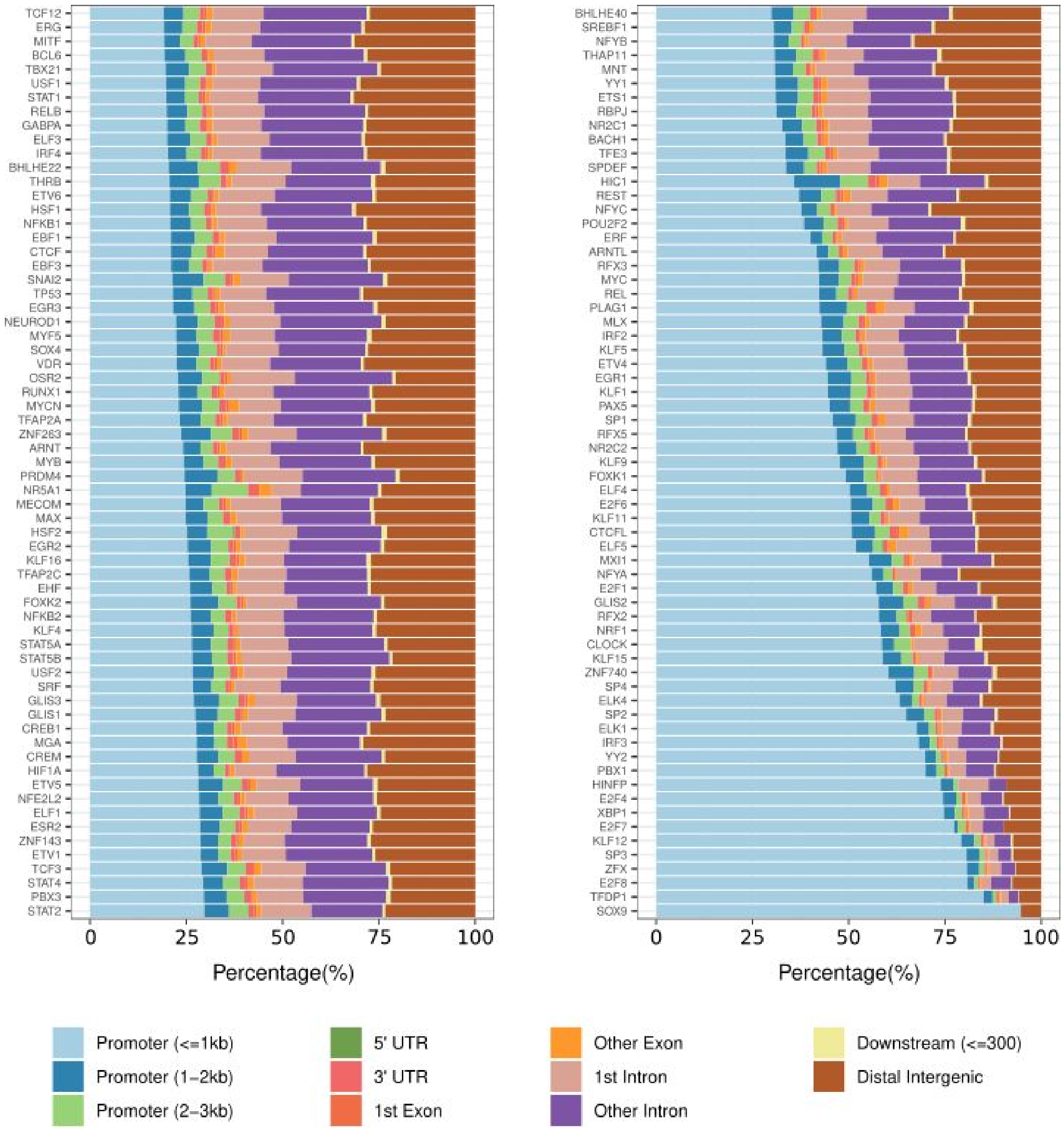

**Supplementary Figure 22 (continued). Genomic distribution of *H. sapiens* TFBSs**. Distribution of the proportion of *H. sapiens* UniBind robust TFBSs overlapping with different types of genomic regions (columns; see legend) across TFs (rows).

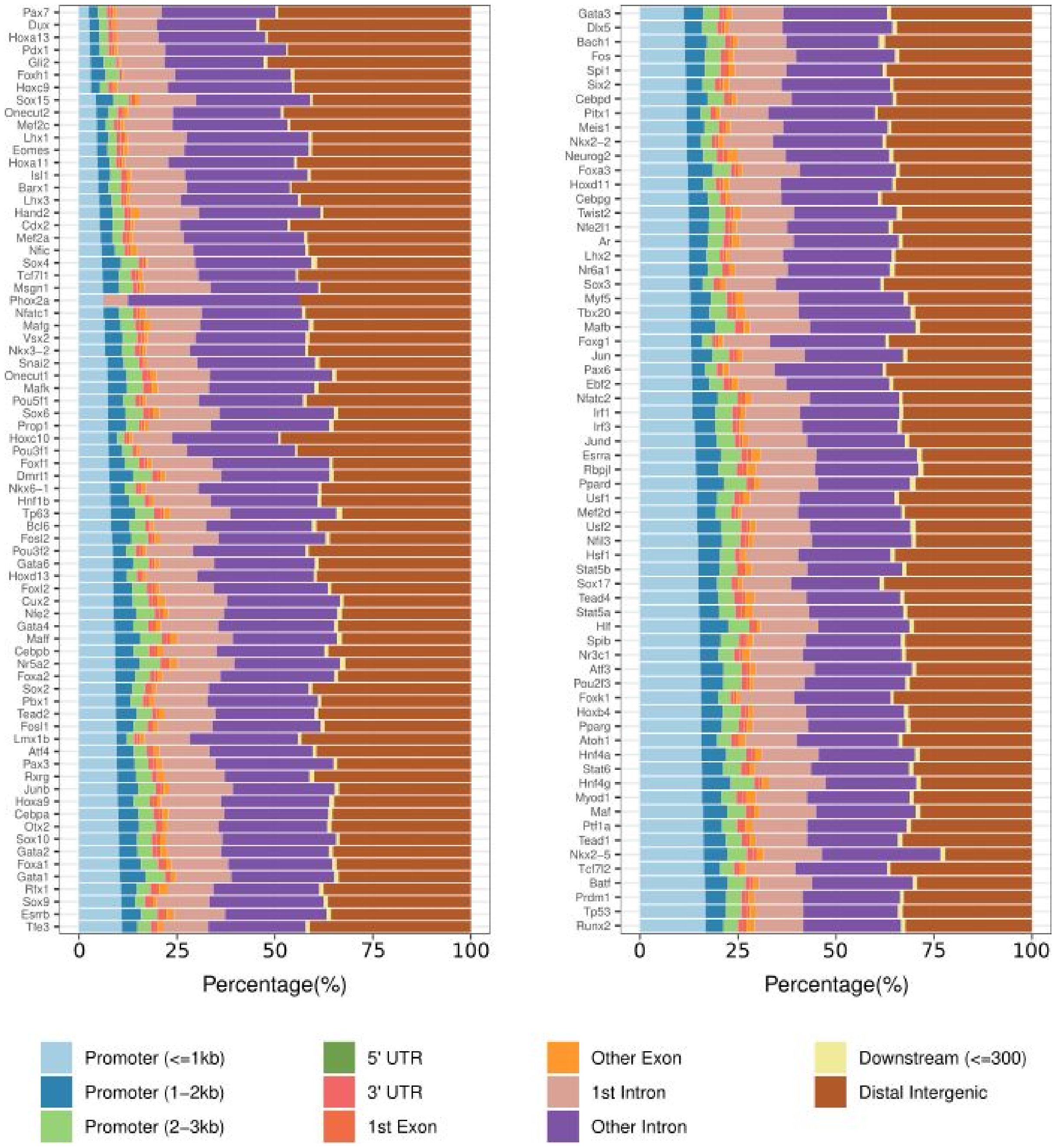


**Supplementary Figure 23. Genomic distribution of *M. musculus* TFBSs**. Distribution of the proportion of *M. musculus* UniBind robust TFBSs overlapping with different types of genomic regions (columns; see legend) across TFs (rows).

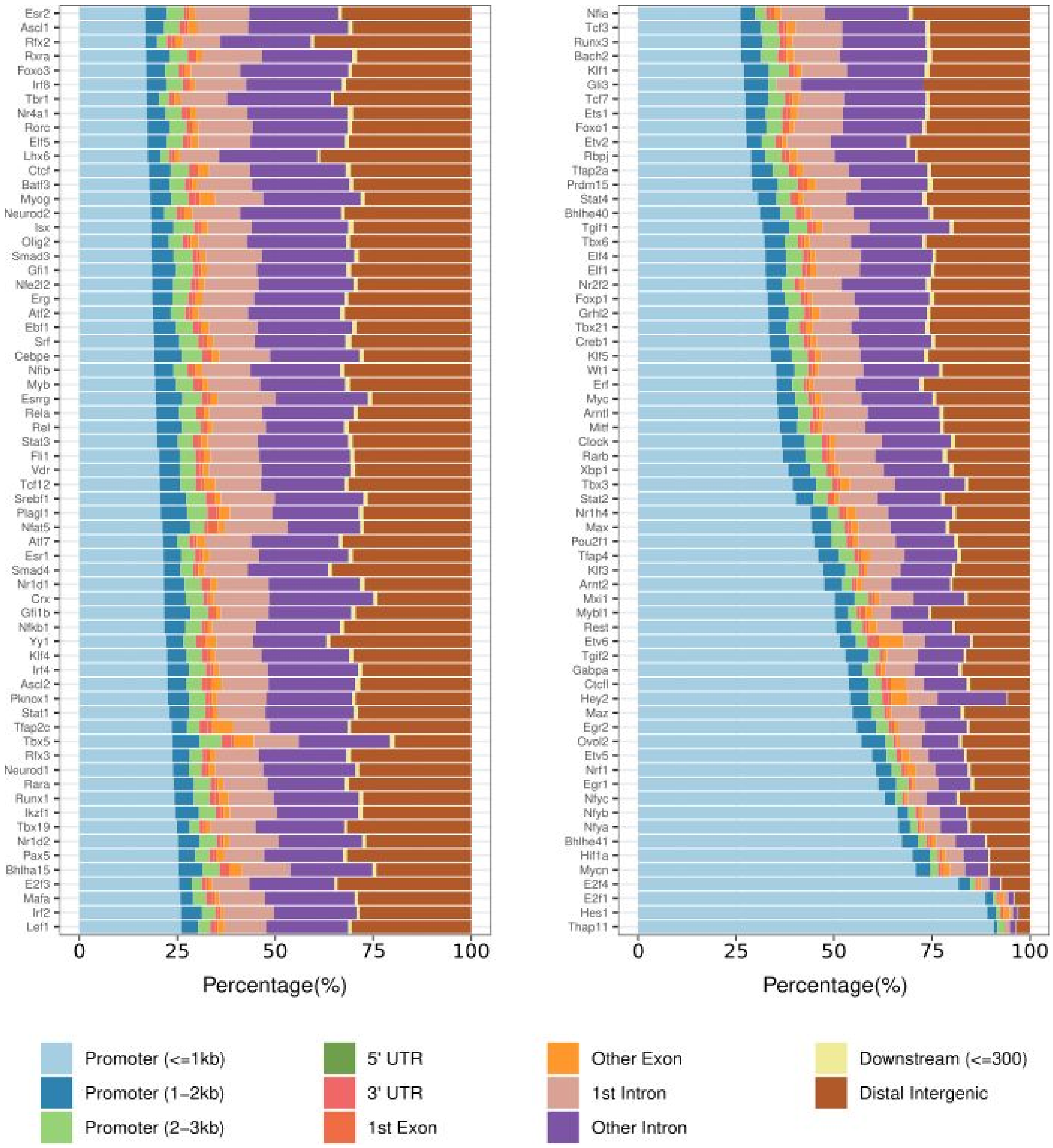


**Supplementary Figure 23 (continued). Genomic distribution of *M. musculus* TFBSs**. Distribution of the proportion of *M. musculus* UniBind robust TFBSs overlapping with different types of genomic regions (columns; see legend) across TFs (rows).

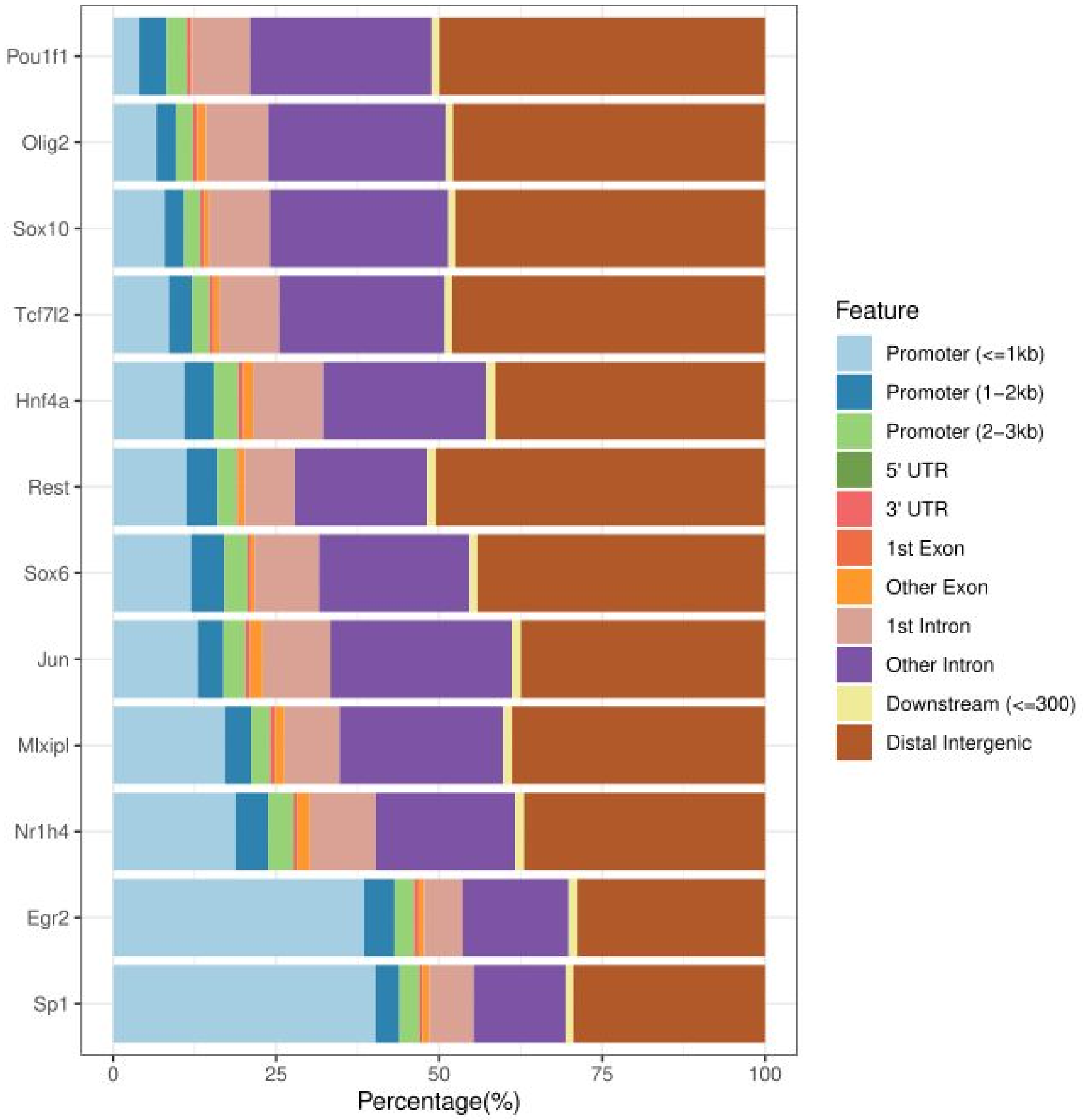


**Supplementary Figure 24. Genomic distribution of *R. norvegicus* TFBSs**. Distribution of the proportion of *R. norvegicus* UniBind robust TFBSs overlapping with different types of genomic regions (columns; see legend) across TFs (rows).

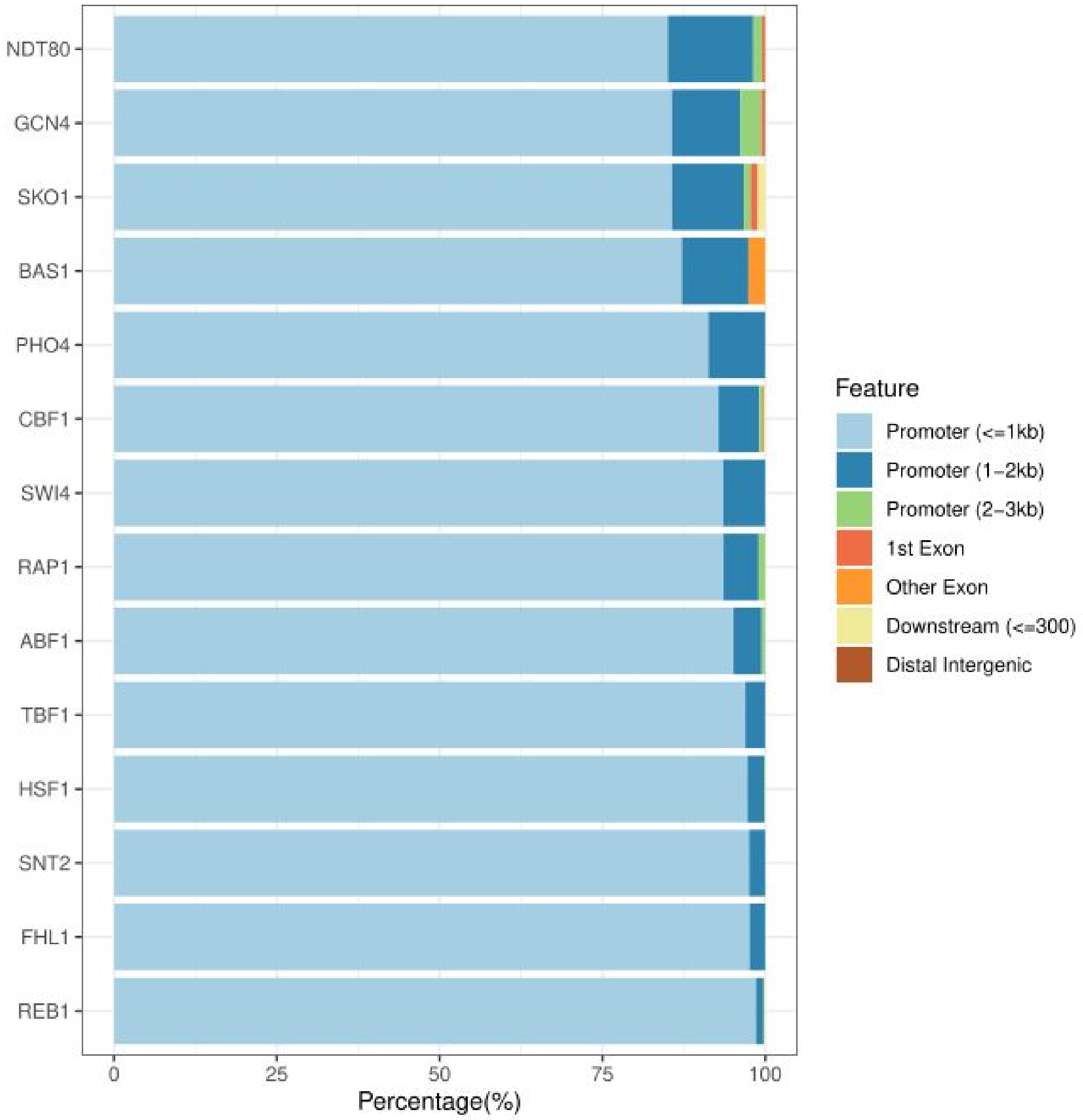


**Supplementary Figure 25. Genomic distribution of *S. cerevisiae* TFBSs**. Distribution of the proportion of *S. cerevisae* UniBind robust TFBSs overlapping with different types of genomic regions (columns; see legend) across TFs (rows).

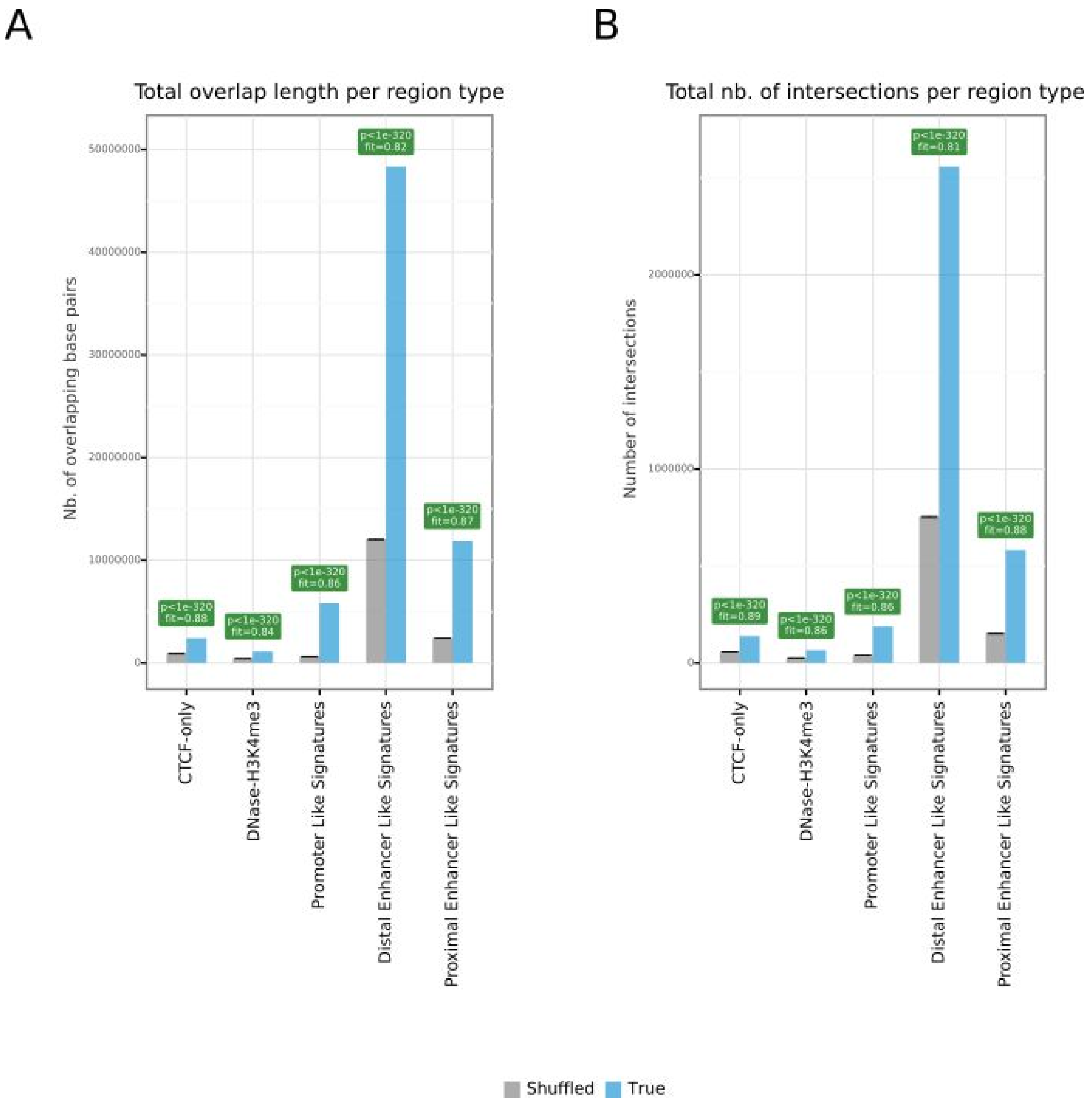


**Supplementary Figure 26**. **Enrichment analysis for *H. sapiens* TFBSs in ENCODE cCREs**. Barplots representing the expected (grey bars) versus observed (blue bars) overlap lengths (**A**) or number of intersections (**B**) between *H. sapiens* TFBSs from the robust collection and ENCODE cCREs (x-axis). The plots and computed p-values (green: enrichment; orange: depletion) were obtained using the OLOGRAM command of the GTF toolkit.

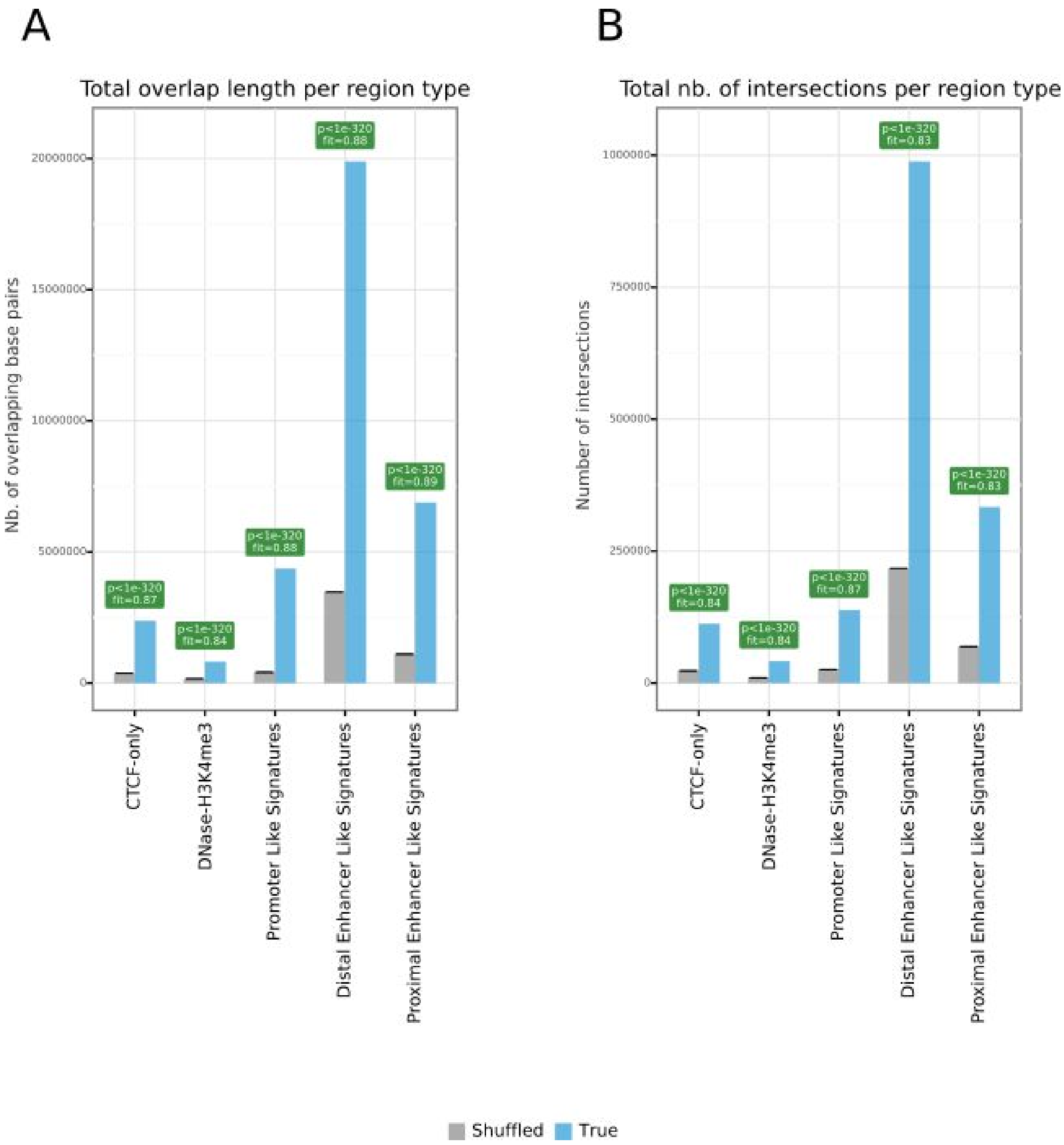


**Supplementary Figure 27**. **Enrichment analysis for *M. musculus* TFBSs in ENCODE cCREs**. Barplots representing the expected (grey bars) versus observed (blue bars) overlap lengths (**A**) or number of intersections (**B**) between *M. musculus* TFBSs from the robust collection and ENCODE cCREs (x-axis). The plots and computed p-values (green: enrichment; orange: depletion) were obtained using the OLOGRAM command of the GTF toolkit.

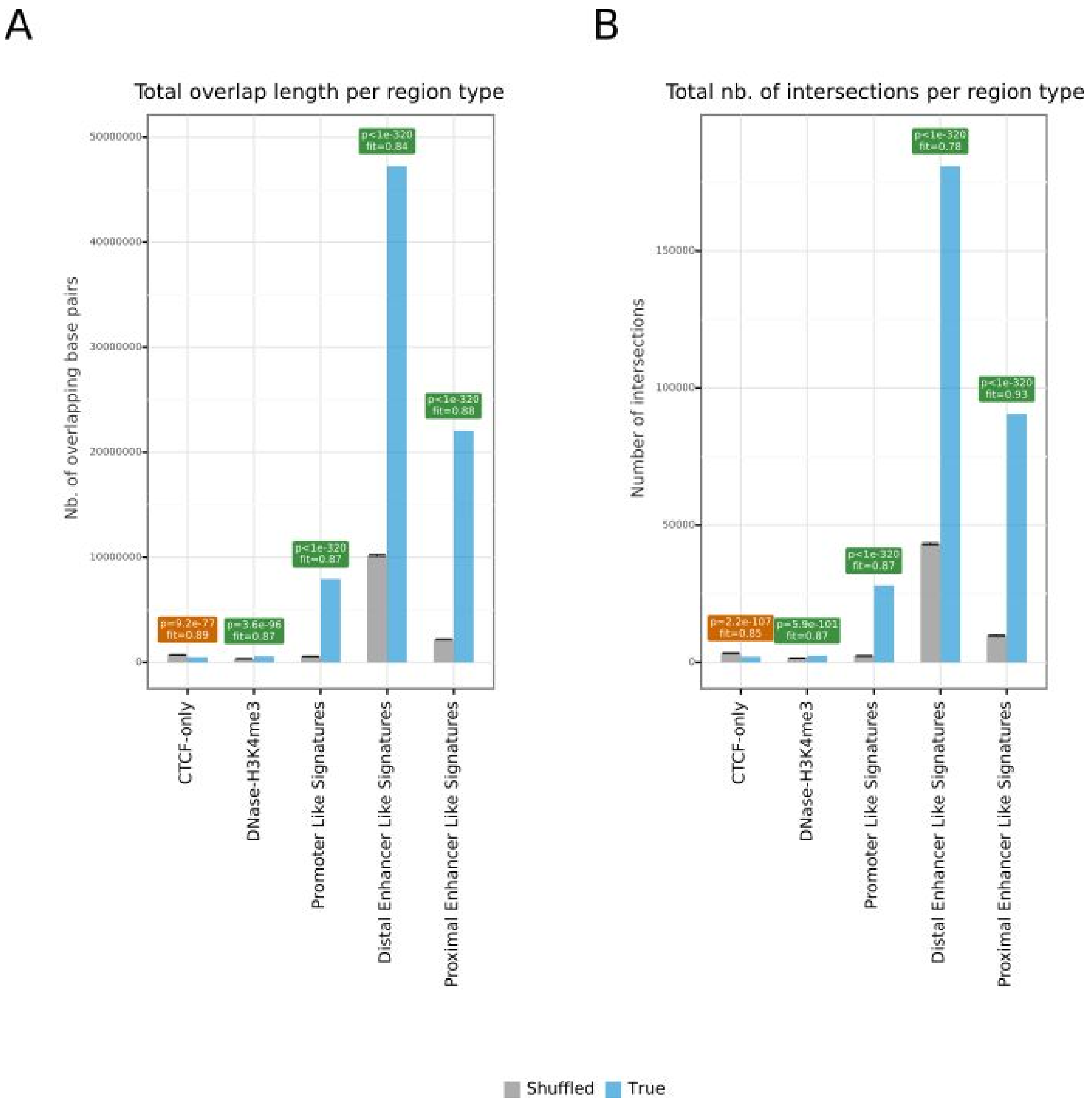


**Supplementary Figure 28**. **Enrichment analysis for *H. sapiens* CRMs in ENCODE cCREs**. Barplots representing the expected (grey bars) versus observed (blue bars) overlap lengths (**A**) or number of intersections (**B**) between *H. sapiens* CRMs from the robust collection and ENCODE cCREs (x-axis). The plots and computed p-values (green: enrichment; orange: depletion) were obtained using the OLOGRAM command of the GTF toolkit.

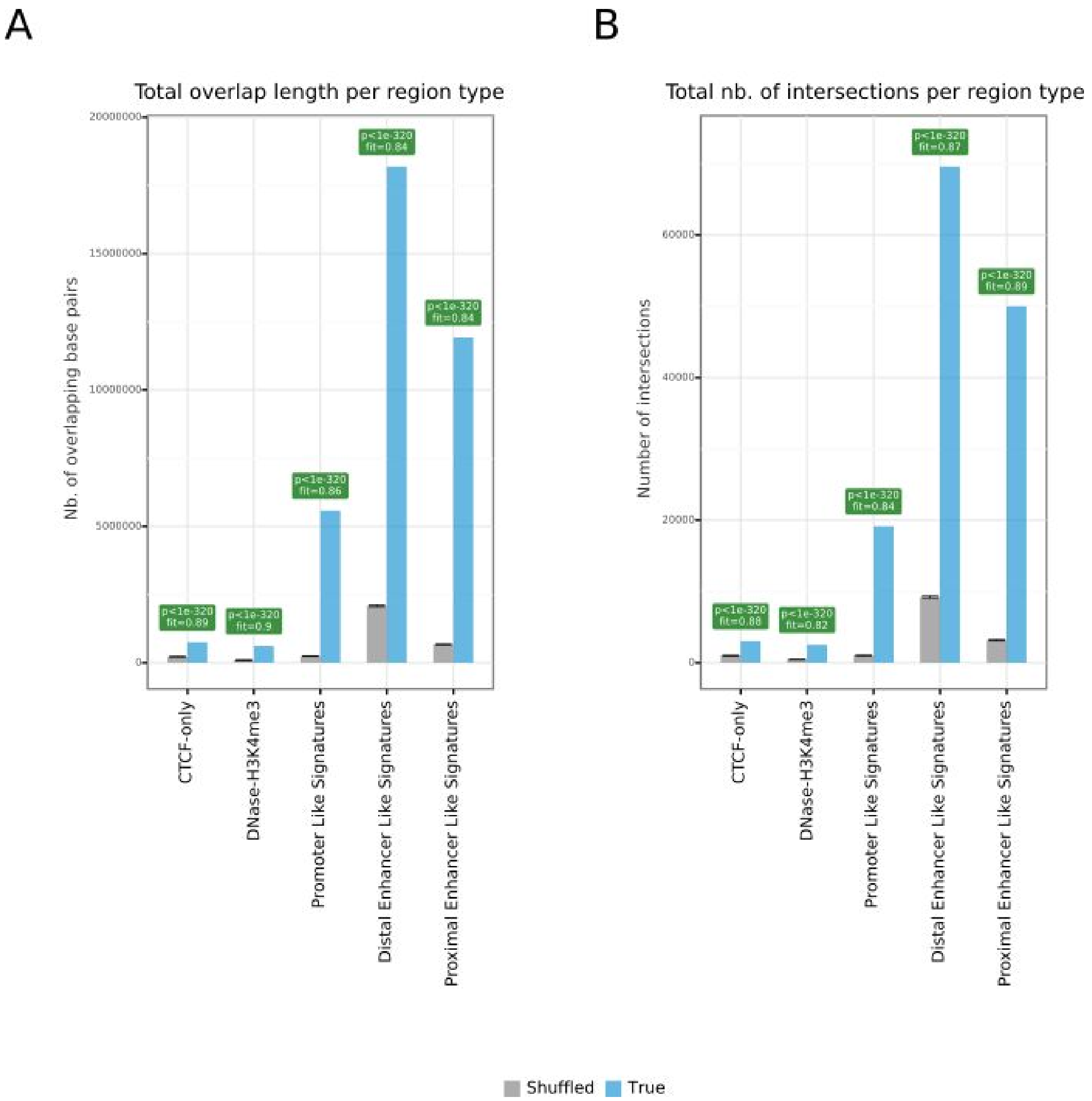


**Supplementary Figure 29**. **Enrichment analysis for *M. musculus* CRMs in ENCODE cCREs**. Barplots representing the expected (grey bars) versus observed (blue bars) overlap lengths (**A**) or number of intersections (**B**) between *M. musculus* CRMs from the robust collection and ENCODE cCREs (x-axis). The plots and computed p-values (green: enrichment; orange: depletion) were obtained using the OLOGRAM command of the GTF toolkit.

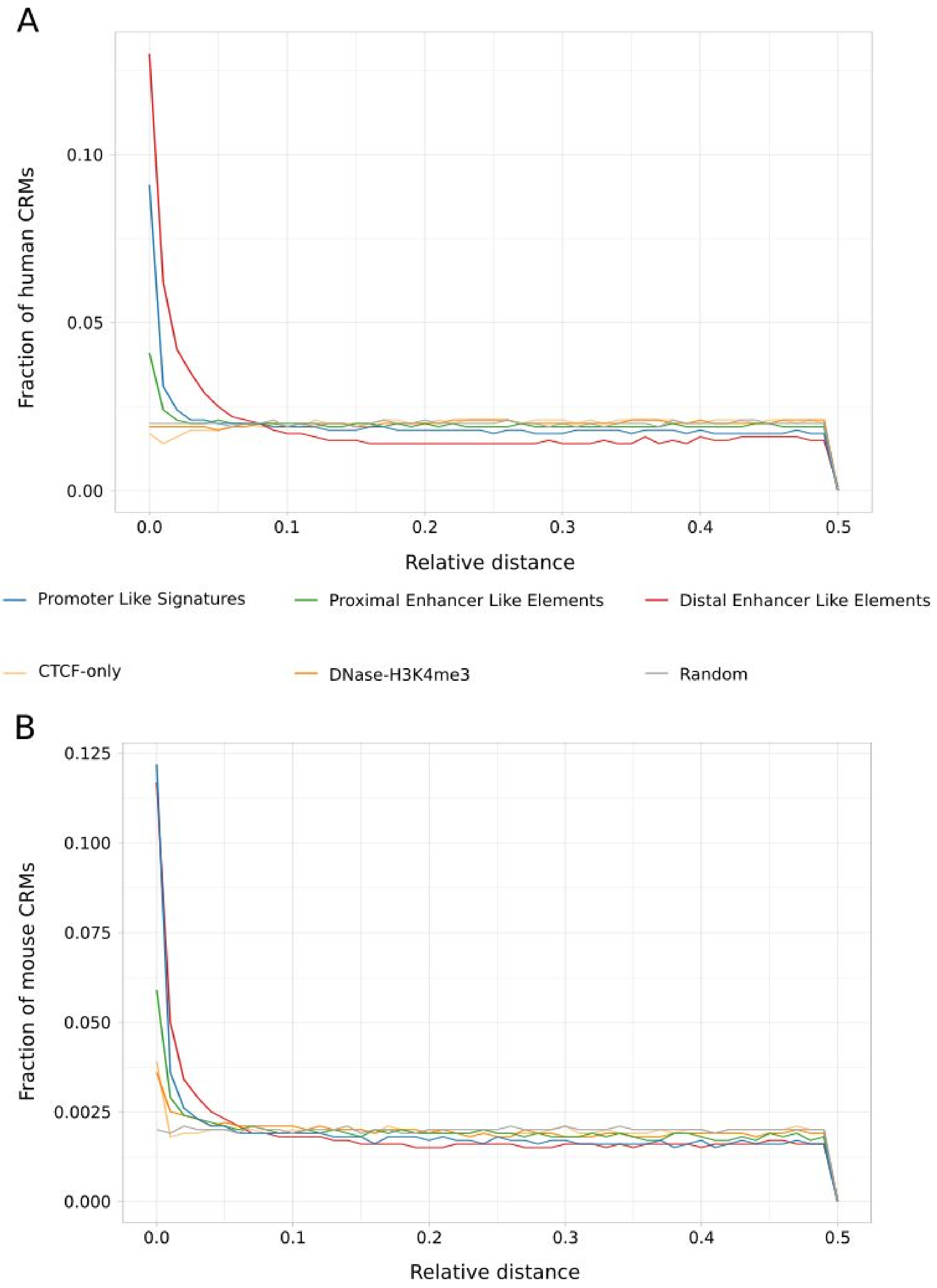


**Supplementary Figure 30. Relative distance distributions between CRMs and ENCODE cCREs.** Fraction of CRMs in the UniBind robust collection (y-axis) with respect to increasing relative distances (x-axis) from ENCODE cCREs computed using the *bedtools reldist* command for human (**A**) and mouse (**B**). When two genomic tracks are not spatially related, one expects the fraction of relative distance distribution to be uniform.

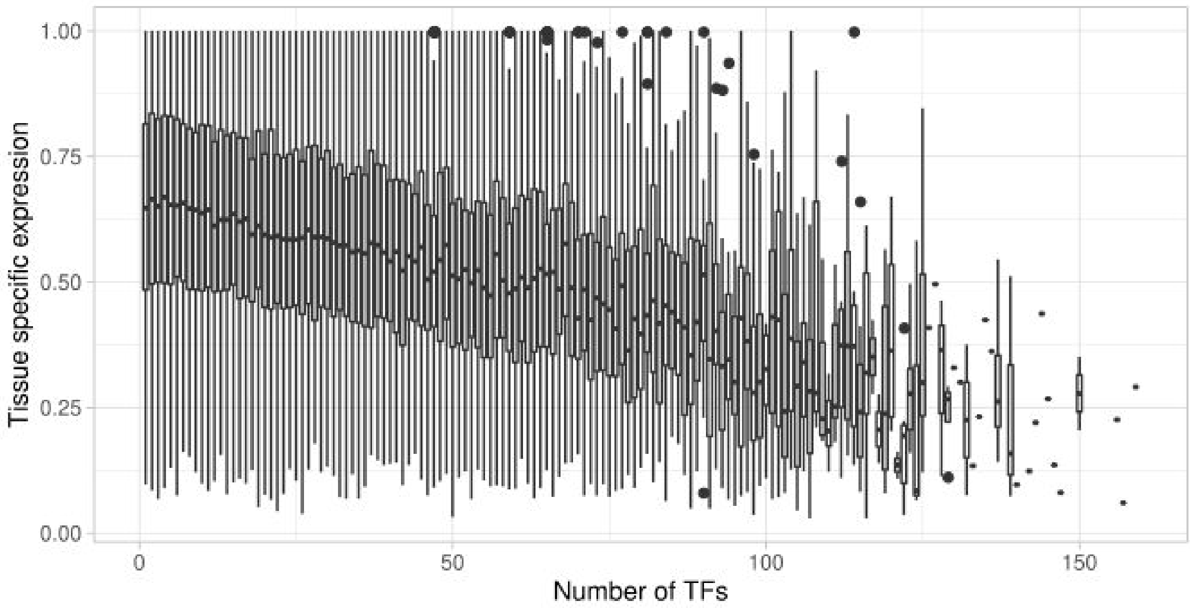


**Supplementary Figure 31**. **Correlation between enhancer activity and TF binding**. For each enhancer predicted using Cap Analysis of Gene Expression (CAGE) by the FANTOM5 consortium, we computed the number of TFs with overlapping TFBSs in the robust collection of UniBind (x-axis). The figure provides, for each value of the number of TFs, a bar plot of the distribution of tissue specific activity of these enhancers. The expression measures were derived from CAGE (capturing enhancer RNA expression). The tissue specificity of activity (y-axis) is provided within the [0; 1] range with 0 representing ubiquitous enhancer activity and 1 exclusive expression activity.
